# Robust self-supervised machine learning for single cell embeddings and annotations

**DOI:** 10.1101/2025.06.05.658097

**Authors:** Christine Yiwen Yeh, Min Woo Sun, Dixian Zhu, Livnat Jerby

## Abstract

Dimensionality reduction and clustering are critical steps in single-cell and spatial genomics studies. Here, we show that existing dimensionality reduction and clustering methods suffer from: (1) overfitting to the dominant patterns while missing unique ones, which impairs the detection and annotation of rare cell types and states, and (2) fitting to technical noise over biological signal. To address this, we developed DR-GEM, a self-supervised meta-algorithm that combines principles in distributionally robust optimization with balanced consensus machine learning. DR-GEM supervises itself by (1) using the reconstruction error to identify and reorient its attention to samples/cells that are otherwise poorly embedded, and (2) using balanced consensus learning as a mechanism to increase robustness and mitigate the impact of low-quality samples/cells. Applied to synthetic and real-world single cell ‘omics data, single cell resolution spatial transcriptomics, and Perturb-seq datasets, DR-GEM markedly and consistently outperforms existing methods in obtaining reliable embeddings, recovering rare cell types, filtering noise, and uncovering the underlying biology. In summary, this study surfaces and addresses a gap in single cell genomics and brings self-supervision to the realm of dimensionality reduction and clustering to better support data-driven discoveries.

## INTRODUCTION

As large volumes of diverse genomics and biomedical data continue to accumulate, dimensionality reduction^1–3^ and clustering^4,5^ have become indispensable tools in the biosciences, as exemplified by applications as disease subtyping and patient risk stratification using clinicogenomic data^6–8^, inference of ancestry in genome-wide association studies (GWAS)^9,10^, and the use of single-cell and spatial genomics to map tissue composition and organization^11–14^, identify novel cell types^15–18^, elucidate molecular mechanisms underlying therapy resistance, and more.

Dimensionality reduction can often adequately represent large multidimensional datasets in a more interpretable lower-dimensional latent space, for example, by condensing correlated features (e.g., genes) into meta-features. This is particularly useful in single-cell genomics, where many genes are highly coexpressed across different cells and can thus be grouped into a single program or signature. While expression levels of individual genes suffer from stochastic measurement noise (e.g., dropouts^19^), aggregating the expression of all genes within a program, this noise (being stochastic) tends to cancel out, resulting in more meaningful representations. These low-dimensional representations of the data are then used for visualization [e.g., t-distributed stochastic neighbor embedding (t-SNE)^20^ or Uniform Manifold Approximation Projection (UMAP)^21^], clustering, and cluster-based annotations (e.g., assigning cells to cell types and subtypes), thus impacting downstream analyses.

Here, we show that, existing dimensionality reduction and clustering pipelines suffer from: (1) overfitting to the dominant patterns while missing unique ones, such that, in the single cell genomics context, the most abundant cells dominate the results and minority/rare cell types and states are left undetected or improperly represented, and (2) fitting to noise and technical artifacts, such that, in the single cell genomics context, doublets or ambient RNA can distort the results.

The first limitation applies to data with latent class imbalance, for example, wherein a distinct population of cells is underrepresented. As we show (**Figure 1, Extended Data Figure 1**), the resulting low-dimensional space learned on such data is prone to poorly embed the minority/rare cells and yield clustering solutions wherein these cells do not form their own cluster and instead get falsely assigned to unrelated clusters. The signal from minority/rare (and often novel and poorly understood) cells is seemingly masked by the more abundant (and often already well-characterized) cells.

**Figure 1:**
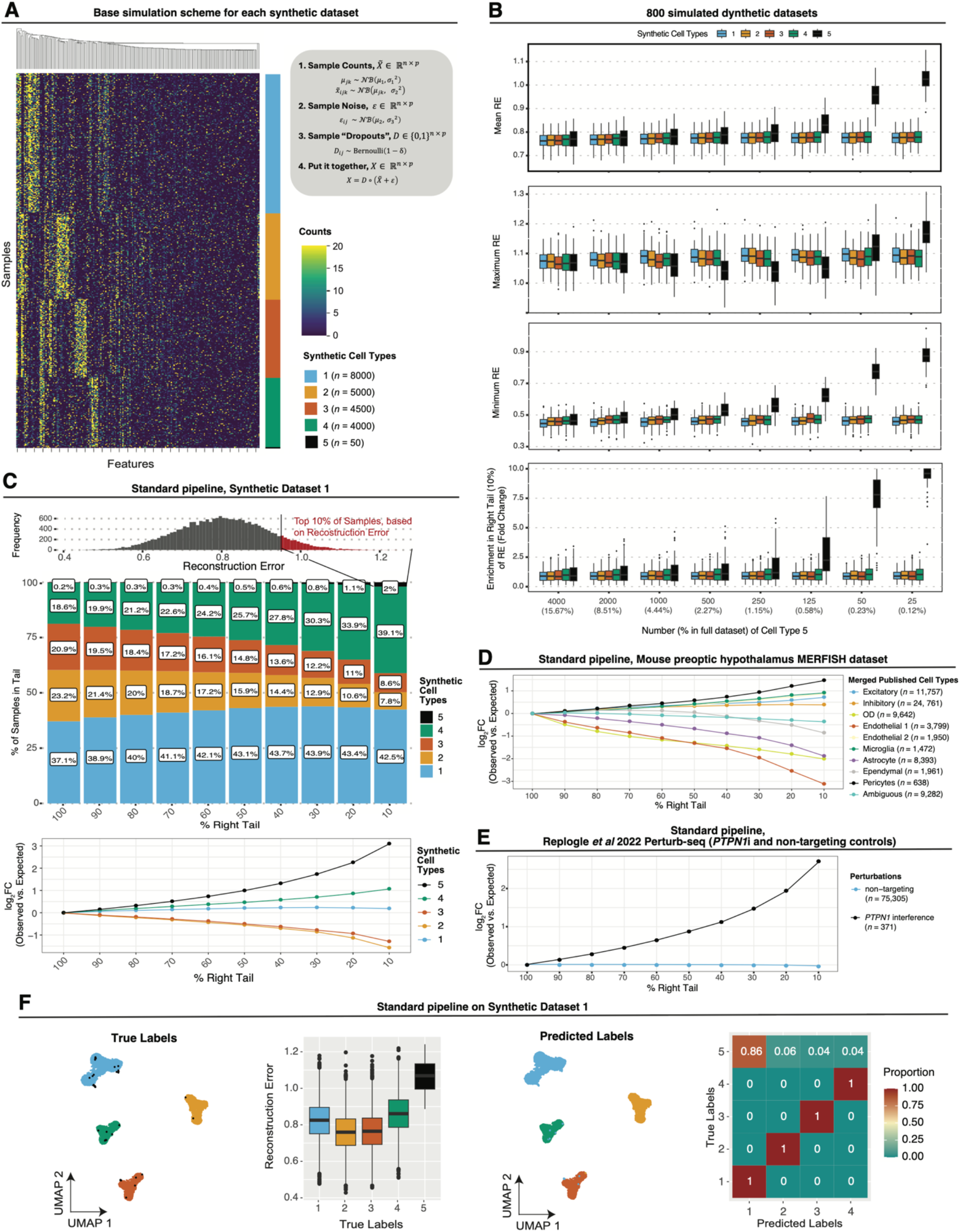
Minority/rare cells are poorly embedded and prone to mislabeling with the standard single cell analysis pipeline. **(A)** Heatmap of Synthetic Dataset 1: values are drawn from two-level negative binomial distributions to model gene expression of 5 synthetic cell types (**Methods**). Data generation is described for cell *i*, gene *j*, and cell type *k*. **(B)** Mean (top), maximum (second row), and minimum (third row) RE, and log_2_FC enrichment in top 10% tail of RE distribution (bottom) of each simulated cell type (fill color) in 100 simulated datasets for each frequency of the minority/rare cell type (*x*-axis). **(C)** Top: histogram of RE in Synthetic Dataset 1. Middle: percentage of each sample population (*y*-axis) in the right-tail (of varying sizes, *x*-axis) in the RE distribution. Bottom: log_2_FC (*y*-axis) of observed percentage of each sample population vs. expected percentage of each sample population in the right-tail (of varying sizes, *x*-axis) in the RE distribution in Synthetic Dataset 1. **(D-E)** log_2_FC (*y*-axis) of observed percentage of each sample population vs. expected percentage of each cell type population in the right-tail (of varying sizes, *x*-axis) in the RE distribution in a mouse preoptic hypothalamus MERFISH dataset^14^ **(D)**, and a Perturb-seq dataset^47^ with non-targeting control and *PTPN1* interference (*PTPN1*i) cells **(E). (F)** Performance of the standard pipeline (**Methods**) on Synthetic Dataset 1: UMAP visualization of samples colored by ground truth synthetic cell type labels (left) and predicted cluster labels (middle left), boxplots of RE (*y*-axis) in each ground truth synthetic cell type (*x*-axis, middle right), and proportion of predicted cell type (*x*-axis) in each of the true cell type populations (*y*-axis; right). Label colors the same as in **(A** and **C)**. Boxplots middle line: median; box edges: 25^th^ and 75^th^ percentiles; whiskers: most extreme points that do not exceed ± interquartile range (IQR) x 1.5; further outliers are marked individually as black points.

Analogous limitations have been observed in the broader context of machine learning and artificial intelligence, where models have been shown to amplify inequities^22–24^ arising from sampling bias and class imbalance^25,26^. Current state-of-the-art solutions amount to either reweighting the data^27,28^ or applying mathematical penalizations of unfair model behavior during model training^29–35^. These principles in algorithmic fairness and distributionally robust optimization (DRO) have been mostly applied to supervised machine learning, and received less attention in the context of unsupervised learning and workflows that include multiple machine learning algorithms, where bias may propagate throughout a pipeline^36–38^.

Other approaches to address latent class imbalance have been focused on data acquisition rather than data analysis. In the context of single cell genomics for example, there have been substantial efforts to discover new (and often rare) cell types and new cell states^15–18^ and comprehensively map all cell types and states across diverse conditions and tissues^15,17,18,39^. Solutions to circumvent poor representation of rare cells have largely been focused on data acquisition, primarily by (1) sequencing more cells^40,41^, or (2) enriching for rare populations of cells^42–46^. While these solutions can partially mitigate some aspects of the problem, they are expensive, cannot enrich for rare cells unless they have already been thoroughly characterized, and do not address limitations that, as we show here, occur in the data analysis stage.

The second limitation of fitting to noise can occur also when the data is of high quality overall, but a fraction of the samples/cells still have a low signal to noise ratio and skew the resulting embedding and clustering. For example, in single cell RNA sequencing (scRNA-Seq), mRNA molecules that have been released in the cell suspension or during tissue dissociation (e.g., from stressed or dying cells) can contaminate the microfluidics droplets and get falsely assigned to other cells. Microfluidic droplets can also sometimes include more than one cell, hence harboring signals that are a mixture of two or more cells^58–60^. In spatial transcriptomics RNA molecules from one cell can get falsely assigned to a nearby cell due to lateral RNA diffusion and limitations of image segmentation^11,61–63^. While improvement in tissue dissociation protocols, microfluidics, and segmentation algorithms help mitigate these problems, these issues are difficult to eliminate completely, calling for computational frameworks that could identify and reorient the model attention to high quality cells to generate embeddings, clustering, annotations, and reference maps with higher quality.

To address these gaps, we developed DR-GEM (**<u>D</u>**istributionally **<u>R</u>**obust and latent **<u>G</u>**roup-Awar**<u>E</u>** consensus **<u>M</u>**achine learning) – a self-supervised meta-algorithm that brings forward and implements the concepts of distributional robustness, data balancing, and consensus learning. DR-GEM outperforms existing unsupervised methods in recovering ground truth information in a range of metrices and prediction tasks, faithfully reconstructing latent transcriptional structure in the context of latent-class imbalance and technical noise, and detecting rare cell populations that are otherwise completely missed by existing methods, as we show using synthetic and real-world single cell, spatial, and perturbational datasets across a variety of technological platforms and biological settings spanning patient tubo-ovarian cancer tumors^11^, mouse preoptic hypothalamus^14^, and CRISPR perturbations in cell models^47^.

## RESULTS

### Minority/rare cells are poorly embedded and prone to mislabeling

Single cell data is often unlabeled at the cellular level and its labeling depends on dimensionality reduction and clustering to assign cells to different cell types, subtypes, and states^7–20^. A canonicalized framework for performing this task strings together several unsupervised algorithms: (1) dimensionality reduction (i.e. principal components analysis (PCA)), (2) clustering (i.e. shared Nearest Neighbors clustering^48^), (3) visualization (e.g., via UMAP two-dimensional embeddings), and (4) annotating clusters as different cell (sub)types or states. While this widely used pipeline is powerful (referred to as the “standard pipeline” hereinafter), we show that it suffers from an intrinsic limitation of poorly embedding and mislabeling minority/rare cell types/states. The quality of the embedding of a sample/cell in this context can be quantitatively determined as higher than expected reconstruction error (RE), that is, the distance between the input data and the (high dimensional) data that is reconstructed from the low dimensional embedding (**Methods**). Mislabeling is defined here as the deviation of the cluster-based labeling from ground truth labels.

To rigorously and quantitatively test and demonstrate the challenges posed by latent class imbalance in a controlled manner that avoids confounders, we simulated synthetic single cell ‘omics data featuring latent class imbalance with ground truth labels (**Figure 1A**). The synthetic data consists of simulated gene expression counts sampled from negative binomial distributions, such that each cell is assigned to one out of *K* cell types. In brief, each of the *k* cell types has a unique set of overexpressed marker genes (**Figure 1A, Methods**; total of *p1* marker genes across all *k* cell types). To more closely simulate real-world single cell ‘omics data, an additional set of *p*_2_ features (where *p*_*2*_ < *p*_*1*_) is added to model a secondary module of cell variation that is invariant to the synthetic cell types. We then add negative binomial noise *ε* and simulate random dropouts by randomly setting a fraction of gene expression count values to 0 (**Figure 1A, Methods**).

Using the scheme described above, we simulated 100 unique synthetic single cell ‘omics datasets for each of 8 different levels of frequency of one minority/rare cell type (Synthetic Cell Type 5) and fit PCA on all 800 simulated datasets. Based on per-cell RE and the ground truth cell type annotations, cells representing the minority/rare cell type are prone to have significantly poorer embeddings as a function of the degree of underrepresentation (**Figures 1B-C**). Examining this in multiple real-world single cell datasets^11,13,14,47,49,50^, minority/rare cells (i.e., the cells of the rare cell type) are 2.85-9.18 times more likely to be among the most poorly embedded cells (cells with the top 10^th^ percentile of RE values); (**Figure 1D-E, Extended Data Figure 1A-I, Methods**).

Subsequent shared nearest neighbor (SNN) clustering on the latent lower dimensional space (**Figure 1F**, *p*= 3.96 × 10^−20^, RE of Synthetic Cell Type 5 vs. Synthetic Cell Type 1-4, one-sided Wilcoxon ranksum test) yielded incorrect cluster assignments for the minority/rare cells in a representative baseline synthetic dataset (Synthetic Dataset 1, **Figure 1F**). The minority/rare cells do not form a cluster of their own and instead are randomly misassigned to other clusters (**Figure 1F**). As we show, this poor embedding (**Figure 1B-F, Extended Data Figure 1A-I)** and erroneous labeling of minority/rare cells also appear in variations on the synthetic data and across several real-world single-cell datasets.

### Existing DRO and fairness-aware algorithms are insufficient to improve the embedding and labeling of minority/rare cells

First, we postulated that the poor embedding and mislabeling of minority/rare samples (e.g., cells) may arise from intrinsic biases with standard error minimization during model fitting for dimensionality reduction. As is true with most machine learning algorithms, the objective of dimensionality reduction is to (either explicitly or implicitly) minimize an average error. Consequently, common patterns in the data dominate the model fitting process. Minority/rare samples contribute less to the overall average error, so models are inherently biased to underperform on them. Conversely, DRO is a family of optimization approaches that shifts the focus from minimizing average loss to better minimize worst-case loss; this prevents models from neglecting rare but important samples, promoting algorithmic fairness and robustness to distributional shifts.

However, DRO dimensionality reduction algorithms do not alleviate the poor embedding and mislabeling of minority/rare samples. Applied to a representative baseline synthetic dataset (Synthetic Dataset 1) Kullback-Leibler Divergence Constrained DRO (KL-DRO^31^) and Conditional Value-at-Risk^51^ DRO (CVaR-DRO) (**Figure 2A**) successfully achieve the DRO objectives to better minimizing worse-case RE for dimensionality reduction across multiple hyperparameters (**Figure 2B**), but do not sufficiently improve the embedding and assignment of rare cells compared to the standard pipeline (**Figures 2C-D**). The minority/rare cells (Synthetic Cell Type 5) continue to have a significantly higher RE compared to all other cell types (*p* = 2.52 × 10^−6^, *p* = 8.98 × 10^−6^ for KL-DRO (λ = 0.001) and CVaR-DO (β = 0.005), respectively, one-sided Wilcoxon ranksum test), do not cluster apart from the other cells, and are mislabeled (**Figure 2C-D**). The lack of improvement in performance with DRO may be explained by the fact that the optimization schemes do not explicitly minimize a single group’s error.

**Figure 2:**
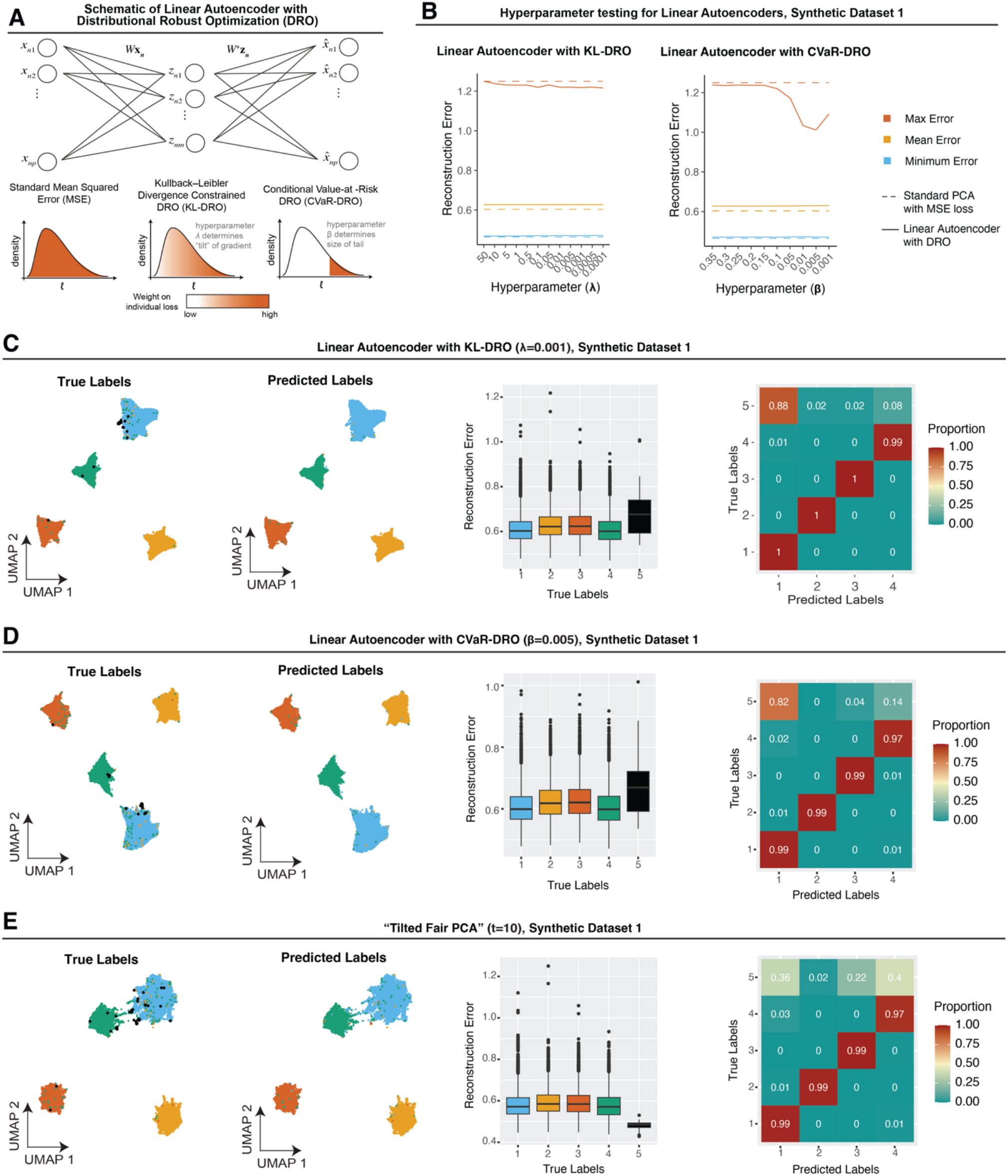
Existing DRO and fairness aware algorithms are insufficient to improve the embedding and labeling of minority/rare cells. **(A)** Schematic of PCA optimized under the KL-DRO^29,31,85^ and CVaR-DRO^51^ objectives (**Methods**). **(B)** Maximum (red), mean (yellow), and minimum (blue) RE (*y*-axis) of the dataset as a function of hyperparameter λ (*x*-axis) in PCA solved via KL-DRO (solid line, left) or CAaR-DRO (solid line, right) vs. default PCA (dotted line). **(C-E)** Example performance of pipeline with PCA solved via KL-DRO **(C)**, CVaR-DRO **(D)**, and “Tilted Fair PCA”^30,52–54^ **(E)** on Synthetic Dataset 1. UMAP visualization of cells colored by ground truth cell type labels (left) and predicted cluster labels (middle left), colors the same as in **Figure 1A**. Middle right: boxplots of RE (*y*-axis) in each ground truth cell type population (*x*-axis). Right: Confusion matrix of true (*y*-axis) vs. predicted (*x*-axis) cell types, each row normalized to sample size of each ground truth cell type. Boxplots middle line: median; box edges: 25^th^ and 75^th^ percentiles; whiskers: most extreme points that do not exceed ± interquartile range (IQR) x 1.5; further outliers are marked individually as black points.

Therefore, we turned to a different conceptual approach to fit “fair” dimensionality reduction (“Tilted Fair PCA”^30,52–54^) wherein the optimization utilizes class labels explicitly. However, even when Tilted FairPCA is given the latent class information (information it would not have access to in real-world setting) and is applied with an optimized hyper-parameter (*t* = 10, **Methods**) to significantly decrease the RE of the rare cell type (**Figure 2E**, *p* = 1, RE of Synthetic Cell Type 5 vs. all other cells, one-sided Wilcoxon ranksum test), the rare cells still do not cluster apart from the other cells and are mislabeled (**Figure 2E**).

These investigations of DRO (KL-DRO and CVaR-DRO) and algorithmic fairness (Tilted Fair PCA) show that existing methods and concepts to mitigate bias in dimensionality reduction are not sufficient to obtain reliable low-dimensional embeddings and accurate labeling for imbalanced datasets. This prompted us to develop an end-to-end self-supervised method that pairs DRO with balanced consensus learning to mitigate these problems, focusing on single-cell analysis.

### DR-GEM is a self-supervised learning algorithm to obtain robust embeddings and labeling of imbalanced datasets

To address the limitations of existing methods, we developed DR-GEM – an iterative self-supervised framework for robust and reliable low-dimensional embeddings and clustering of datasets with latent class imbalance.

DR-GEM has three main Phases. In Phase I (Distributional Robust Initialization, **Figure 3A**), it first fits a standard embedding (*E*^(0)^) using classic dimensionality reduction (e.g., PCA) and computes the RE per cell (**Figure 3A**, left). Under the key observation that minority/rare latent populations tend to be enriched in the right tail of the RE distribution, DR-GEM fits a new embedding *E*^(init)^ on cells with the highest RE values from *E*^(0)^ (**Figure 3A**, middle), thereby shifting the dimensionality reduction algorithm’s focus to these putative high-risk cells. DR-GEM then projects all remainder cells to the *E*^(init)^ subspace and uses this embedding of cells to identify initial clusters ***c***^(1)^ (**Figure 3A**, right).

**Figure 3:**
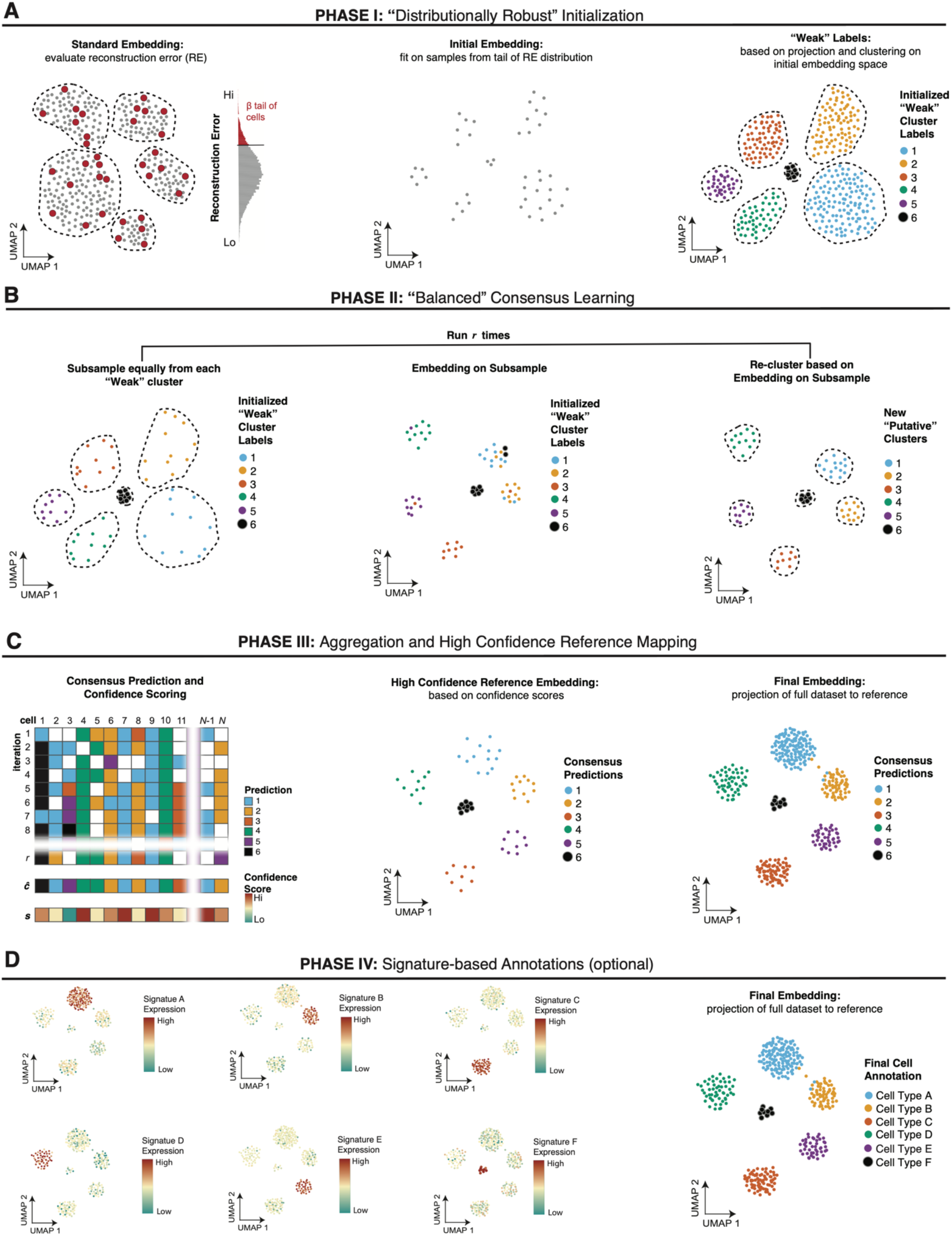
DR-GEM is a self-supervised meta-algorithm for dimensionality reduction and clustering. **(A)** Phase I: Standard dimensionality reduction is performed on the entire dataset to compute RE per cell (left); cells in the *β* right tail of the RE distribution are used to generate an initial embedding (*E*^(init)^, middle); all cells in the dataset are projected to *E*^(init)^ and clustered to obtain “weak” clustering labels (***c***^(1)^). **(B)** Phase II: *r* unique, random, but balanced (across ***c***^(1)^ “weak” cluster labels) subsamples of the dataset are separately embedded and clustered to generate *r* sets of latent class predictions. **(C)** Phase III: Aggregation of ensemble predictions across iterations from (B) yields consensus cluster labels (**ĉ**) and sample label confidence (***s***) (left); a balanced subsample of “high confidence” cells based on scores ***s*** is used to generate a new “high confidence” reference embedding (middle) to which all cells are projected for a fairness-aware visualization (right). **(D)** DR-GEM includes optional functionality to utilize the expression of gene signatures in each cluster (left) to automatically assign clusters to known cell type/states (right). See **Methods** for more detail.

In Phase II (Balanced Consensus Learning; **Figure 3B**), DR-GEM applies consensus learning, a technique in ensemble machine learning where an array of models is fit, each based on a subset of the data, and the results can then be aggregated to increase accuracy and robustness. To promote latent class balance, DR-GEM randomly samples uniformly from the initial clusters ***c***^(1)^ (**Figure 3B, Methods**), effectively upweighting minority/rare cells across each of the *r* unique subsamples. DR-GEM fits a lower-dimensional embedding and generates cluster annotations for each of the *r* unique subsamples. This yields an ensemble of cluster predictions for each cell. Cluster annotations in each run are aligned via the Kuhn-Munkres^55,56^ algorithm to consolidate the annotations (**Methods**).

In Phase III (Aggregation and High Confidence Reference Mapping; **Figure 3C**), DR-GEM aggregates the solutions by assigning each cell to the cluster it is most frequently assigned to in Phase II. A confidence score is also computed per cell to denote the fraction of runs where the cell was assigned to its final cluster. Lastly, DR-GEM uniformly samples high-confidence cells across the final clusters to generate a final reference embedding to which all the cells are projected for visualization and clustering. High-confidence cells are defined as cells with *s* > *t*, where *t* is a user-defined parameter. As we show, high-confidence cells are less likely to be contaminated with ambient RNA or be doublets, thus generating an embedding that is less impacted by these technical confounders.

In an optional Phase IV (Signature-based Annotation; **Figure 3D**), DR-GEM uses predefined gene signatures of cell types or cell states to automatically assign clusters to known cell types or cell states and support end-to-end single-cell annotations (**Methods**). This step also supports intermediate cell type/state predictions, as we describe in the **Methods** (DR-GEM with Annotations-in-the-loop).

We note that DR-GEM can be used with any dimensionality reduction or clustering algorithm as input. The results shown here were derived by applying PCA and SNN clustering, a standard pair of algorithms used in single-cell analyses.

### DR-GEM successfully recovered latent patterns and labels in synthetic datasets with latent class imbalance and ambient RNA

Applying DR-GEM to Synthetic Dataset 1 shows that, unlike all other pipelines (PCA, KL-DRO, CVaR-DRO, and Fair-PCA; **Figures 1F, 2C-E**), only with DR-GEM the rare cells form their own cluster and are correctly annotated (98%, **Figure 4A**). DR-GEM balances out the RE between the cell groups (*p* = 0.78, RE of Synthetic Cell Type 5 vs. all other cells, one-sided Wilcoxon ranksum test) and improves the prediction of the ground truth cell labels across all balanced clustering prediction performance^57^ metrics (**Figure 4A, Supplementary Table S1A**). As KL-DRO, CVaR-DRO, and Fair-PCA did not outperform the standard pipeline, we next focused on benchmarking DR-GEM’s performance against the standard pipeline with an array of synthetic and real-world datasets.

**Figure 4:**
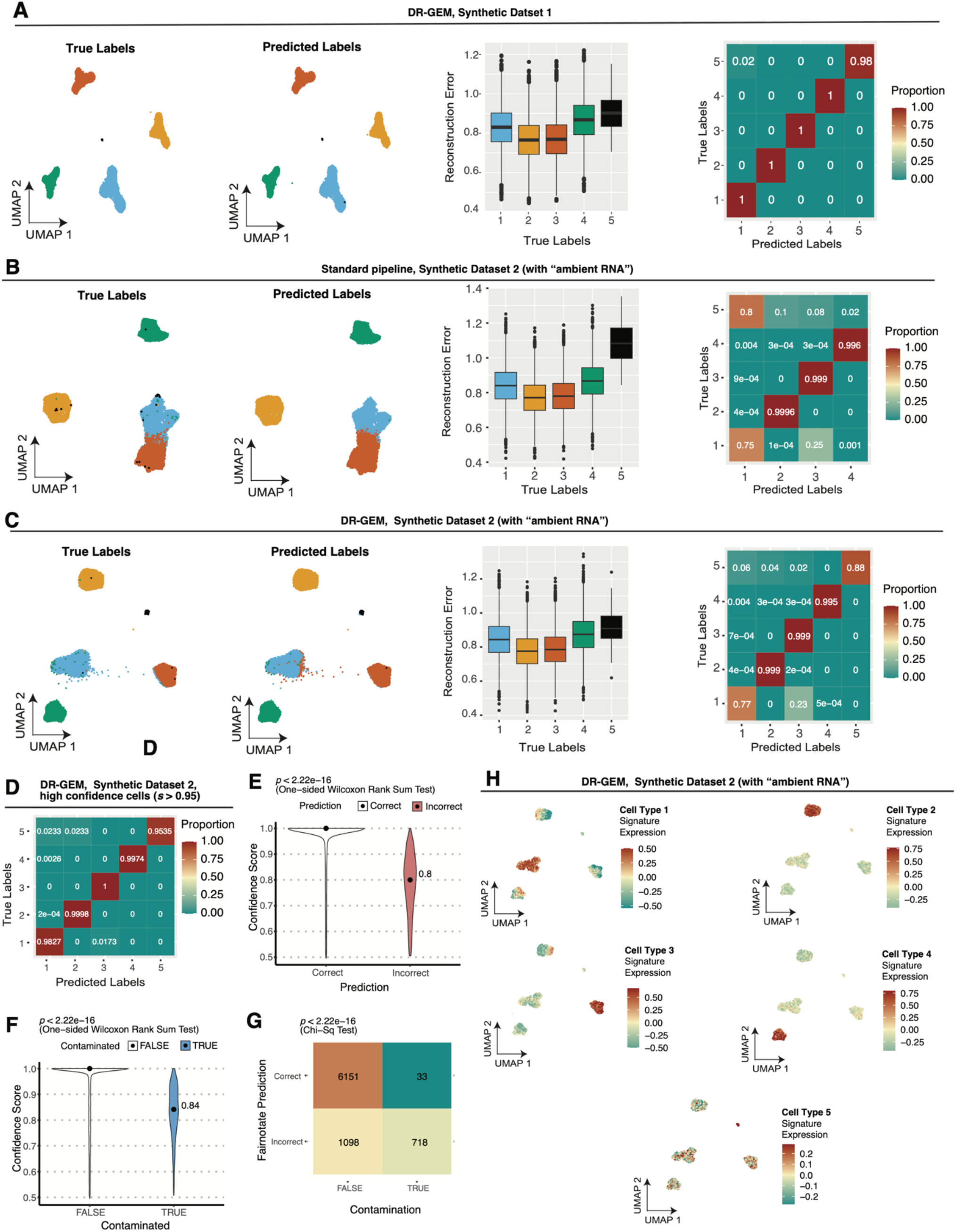
DR-GEM outperforms alternative methods and improves the embedding and labeling of minority/rare cells in synthetic datasets. **(A-C)** Performance of **(A)** DR-GEM on Synthetic Dataset 1 **(B)** standard pipeline on Synthetic Dataset 2 (with “ambient RNA”), and **(C)** DR-GEM on Synthetic Dataset 2. **(A-C)** UMAP visualization of cells colored by ground truth synthetic cell type labels (left) and predicted cluster labels (middle left), boxplots of RE (*y*-axis) in each ground truth cell type population (*x*-axis, middle right), and proportion of predicted cell type (*x*-axis) in each of the true cell type populations (*y*-axis; right). Label colors the same as in **Figure 1A**. Boxplots middle line: median; box edges: 25^th^ and 75^th^ percentiles; whiskers: most extreme points that do not exceed ± interquartile range (IQR) x 1.5; further outliers are marked individually as black points. Violin plots: median denoted with dot labeled with median value. (**D-H**) Additional performance metrics for DR-GEM applied to Synthetic Dataset 2. **(D)** Confusion matrix of confidence-filtered (subset of cells where *s*_*i*_ > 0.95) Synthetic Dataset 2: true (*y*-axis) vs. predicted (*x*-axis) labels with each row normalized to sample size of each true cell type. **(E-F)** Violin plots of DR-GEM confidence scores (*y*-axis) as a function of **(E)** whether a cell was “contaminated” or not (*x*-axis), and **(F)** whether the prediction was correct or not. **(G)** Frequency (color bar) of sample label prediction (*y*-axis) as a function of sample contamination status (*x*-axis). **(H)** UMAP visualization of high confidence cells in the high confidence reference embedding from DR-GEM. Cells colored by overall expression (**Methods**) of multi-feature “signature” of simulated Cell Types 1-5.

Testing DR-GEM in a more challenging setting, we generated Synthetic Dataset 2, which features both simulated imbalanced latent labels and ambient RNA (**Methods**). DR-GEM outperformed the standard pipeline in Synthetic Dataset 2, as it: (1) equalized RE across the synthetic cell types (*p* = 1.40 × 10^−27^, *p* = 0.75 RE of rare vs. all other cell types, one-sided Wilcoxon ranksum test in the standard vs. DR-GEM pipeline, respectively; **Figure 4B-C**), (2) outperformed on all balanced clustering metrics^57^ (**Methods, Supplementary Table S1A**), and (3) clustered the rare cells as a separate cluster, such that 88% of the rare cells were correctly annotated (**Figure 4C**), compared to 0% with the standard pipeline (**Figure 4B**).

DR-GEM assigned 86.9% and 99.0% of the cells with ambient RNA to their correct label when considering all contaminated cells or only those with high confidence scores (*s* > 0.95), respectively (**Figures 4D-H**), while the standard pipeline assigned 84.3% of the cells with ambient RNA to their correct label. Considering only the contaminated rare cells, DR-GEM and the standard pipeline assigned 64.7% and 0% of these cells to the correct label, respectively. The confidence scores correctly identified poorly annotated cells and were significantly lower for cells with incorrect assignments (**Figure 4E**) and contaminated cells (**Figure 4F**).

### DR-GEM outperformed the standard pipeline when applied to spatial transcriptomics datasets profiling tubo-ovarian cancer

To test DR-GEM in end-to-end single cell annotations in a real dataset with documented latent class imbalance and ambient RNA, we applied both the standard pipeline and DR-GEM to a subset of a large-scale single-cell resolution spatial transcriptomics dataset (collected on CosMx Single Molecule Imaging platform (SMI)^11^) profiling non-malignant cells from 47 high grade serous ovarian carcinoma tumors from the omentum from a total of 41 patients (**Methods**).

The cell type annotations in this dataset reported in the original paper were derived from a combination of different methods including data-integration with scRNA-seq, clustering of the entire data, clustering within clusters, matching with tissue histology, and copy number alterations data. We treated these cell type labels as ground truth: Fibroblast (*n*=32,275), Monocyte (*n*=22,791), Endothelial (*n* = 6,081), B cell (*n* = 7,705), T/NK cell (*n* = 13,627), Mast cell (*n* = 329). The Mast cells were the rarest cell type, accounting for 0.397% in the dataset (**Extended Data Figure 1A**).

With the standard pipeline, the Mast cells are scattered across the embedding instead of forming a Mast cell cluster, have significantly higher RE values compared to all other cell types (*p* = 5.08 × 10^−206^, RE of Mast cells vs. all other cell types, one-sided Wilcoxon ranksum test), and are disproportionately enriched among the cells with higher RE values (**Figure 5A**; **Extended Data Figure 1A**). Clustering followed by the Kuhn-Munkres matching algorithm identified one-to-one matching clusters corresponding to Fibroblasts (96% accuracy), Monocytes (97% accuracy), Endothelial cells (98%, accuracy), B cells (61% accuracy), and T/NK cells (97% accuracy), but no cluster corresponded to Mast cells.

**Figure 5:**
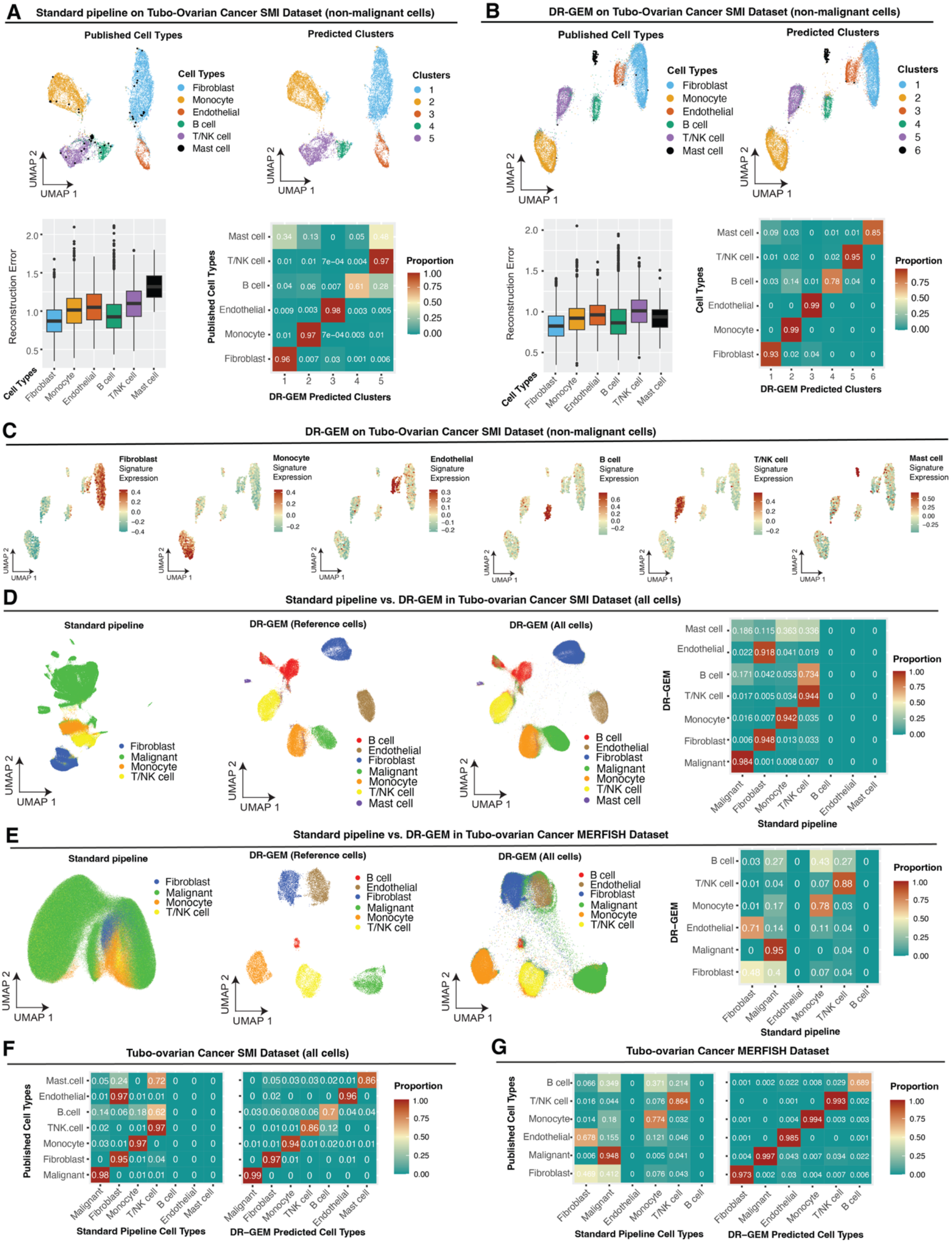
DR-GEM identifies rare cell types and mitigates ambient RNA in spatial transcriptomics datasets profiling tubo-ovarian cancer. **(A-B)** Performance of **(A)** standard pipeline and **(B)** DR-GEM when applied to non-malignant cells in a tubo-ovarian cancer SMI dataset^47^. UMAP visualization of cells colored by published cell type labels (top left) and predicted clusters (top right), boxplots of RE (*y*-axis) in each cell type population (*x*-axis; bottom left), and proportion of predicted cell type (*x*-axis) in each of the published cell type populations (*y*-axis; bottom right). **(C)** UMAP visualization of high confidence cells in the high confidence reference embedding generated by DR-GEM when applied to tubo-ovarian cancer SMI dataset. Cells colored by overall expression of cell-type signatures derived from publicly available tubo-ovarian cancer scRNA-seq datasets (**Methods**). **(D-E)** Performance of standard pipeline vs. DR-GEM when applied to full tubo-ovarian cancer datasets profiled by **(D)** SMI and **(E)** MERFISH. UMAP visualization of cells colored by cell type labels derived from the standard pipeline (left); UMAP visualization of high confidence reference cells colored by cell type labels predicted by DR-GEM (middle left); UMAP visualization of all cells colored by cell type labels predicted by DR-GEM (middle right), proportion of standard pipeline cell type predictions (*x*-axis) in each of the DR-GEM cell type populations (*y*-axis; right). **(F-G)** and proportion of standard pipeline (*x*-axis, right) and DR-GEM (*x*-axis, right) predicted cell types in each of the true cell type populations (*y*-axis) in tubo-ovarian cancer tumors profiled by **(F)** SMI, and **(G)** MERFISH. **(A-B)** Boxplots middle line: median; box edges: 25^th^ and 75^th^ percentiles; whiskers: most extreme points that do not exceed ± interquartile range (IQR) x 1.5; further outliers are marked individually as black points.

In contrast, with DR-GEM, Mast cells form a Mast cell cluster and RE values in this cluster are comparable to those of other cell types (**Figure 5B**, *p* = 0.303, RE of Mast cells vs. all other cell types, one-sided Wilcoxon ranksum test). Clustering under DR-GEM yielded one-to-one matching clusters corresponding to all cell types (**Figure 5B**): Fibroblasts (93% accuracy), Monocytes (99% accuracy), Endothelial cells (99%, accuracy), B cells (78% accuracy), and T/NK cells (95% accuracy), and Mast cells (85% accuracy). DR-GEM also outperforms the standard pipeline on all balanced clustering metrics (**Methods, Supplementary Table S1A**). Using cell type signatures previously derived from a meta-analysis^11^ of high grade serous ovarian carcinoma single-cell RNA-seq datasets, DR-GEM Phase IV correctly mapped the clusters to cell types, perfectly matching the cluster annotations based on the ground truth labels (**Figure 5C, Methods**).

We then applied the standard pipeline and DR-GEM (with “Annotations in the Loop”, **Methods**) on the full single-cell resolution spatial transcriptomics dataset profiling tubo-ovarian cancer tumors collected via SMI^11^ (**Methods**). We treated these labels from the original study as ground truth (listed in order of decreasing abundance): Malignant (*n* = 314,191), Fibroblast (*n* = 72,861), Monocyte (*n* = 45,549), T/NK cell (*n* = 28,676), B cell (*n* = 16,373), Endothelial (*n* = 13,536), Mast cell (*n* = 606). With the standard pipeline, only the top 4 most abundant cell types were detected (**Figure 5D**,**F**): Malignant (98%), Fibroblast (95%), Monocyte (97%), and T/NK cells (97%) (**Figure 5D**). Meanwhile, DR-GEM was able to detect all 7 cell types: Malignant (98%), Fibroblast (97%), Monocyte (94%), and T/NK cell (86%), B cells (70%), Endothelial (96%), and Mast cell (86%) (**Figure 5F**), and generate robust embeddings (**Figure 5D**).

To evaluate DR-GEM’s generalizability across platforms, we tested both the standard pipeline and DR-GEM (with “Annotations in the Loop”, **Methods**) on another full single-cell resolution spatial transcriptomics dataset also profiling tubo-ovarian cancer tumors, but collected via MERFISH^11^ (Multiplexed Error-Robust Fluorescence In Situ Hybridization, **Methods**). We treated these labels from the original study as ground truth (listed in order of decreasing abundance): Malignant (*n* = 334,330), Fibroblast (*n* = 37,775), Monocyte (*n* =35,017), T/NK cell (*n* = 8,238), Endothelial (*n* = 5,890), B cell (*n* =4,328). With the standard pipeline, only the top 4 most abundant cell types were detected (**Figure 5E,G**): Malignant (95%), Fibroblast (47%), Monocyte (77%), and T/NK cells (86%) (**Figure 5D**). Meanwhile, DR-GEM was able to detect all 6 cell types: Malignant (99.7%), Fibroblast (97.3%), Monocyte (99.3%), and T/NK cell (99.3%), and B cells (68.9%), Endothelial (98.5%), (**Figure 5F**) and generate robust embeddings (**Figure 5D**).

### DR-GEM outperformed the standard pipeline when applied to MERFISH spatial transcriptomics data of mouse hypothalamus

DR-GEM outperformed the standard pipeline also when applied to a MERFISH dataset profiling the preoptic hypothalamus of one animal^14^. The cell type annotations in this dataset reported in the original paper were derived from a combination of different methods including data-integration with scRNA-seq, clustering of the entire data, clustering within clusters, and differential expression of specific marker genes, etc. In particular, the labeling of neurons as excitatory vs. inhibitory was based on a few marker genes rather than more global transcriptional differences. As ground truth labels, we used the cell type and subtype labels from the original paper (**Methods**): Excitatory (*n*=11,757), Inhibitory (*n*=24,761), oligodendrocytes (OD, *n*=9,642), Endothelial 1 (*n*=1,950), Endothelial 2 (*n*=1,950), Microglia (*n*=1,472), Astrocyte (*n*=8,393), Ependymal (*n*=1,950), Pericytes (*n=*638). We also included cells that were labeled as “Ambiguous” (*n*=9,382) as an opportunity to test DR-GEM’s ability to identify cells that may be subject to ambient RNA. The rarest cell type in this setting is the Pericyte, representing 0.866% of the full dataset.

With the standard pipeline, Pericytes either cluster with Endothelial 2 cells or are scattered throughout the UMAP embedding (**Figure 6A**). RE of Pericytes is significantly higher compared to all other cell types (*p* = 1.50 × 10^−3^, one-sided Wilcoxon ranksum test), such that Pericytes are disproportionately enriched amongst the most poorly represented cells (**Figure 6A**). There is a wide spread of RE for the cells that were labeled as “Ambiguous” by the authors (**Figure 6A**). Clustering and cluster annotation predicted the following cell types: Excitatory (63% accuracy), Inhibitory (93% accuracy), OD (90%, accuracy), Endothelial 1 (83% accuracy), Endothelial 2 (95% accuracy), Microglia (56% accuracy), Astrocytes (90% accuracy), Ependymal (96% accuracy). None of the clusters corresponded to Pericytes.

**Figure 6:**
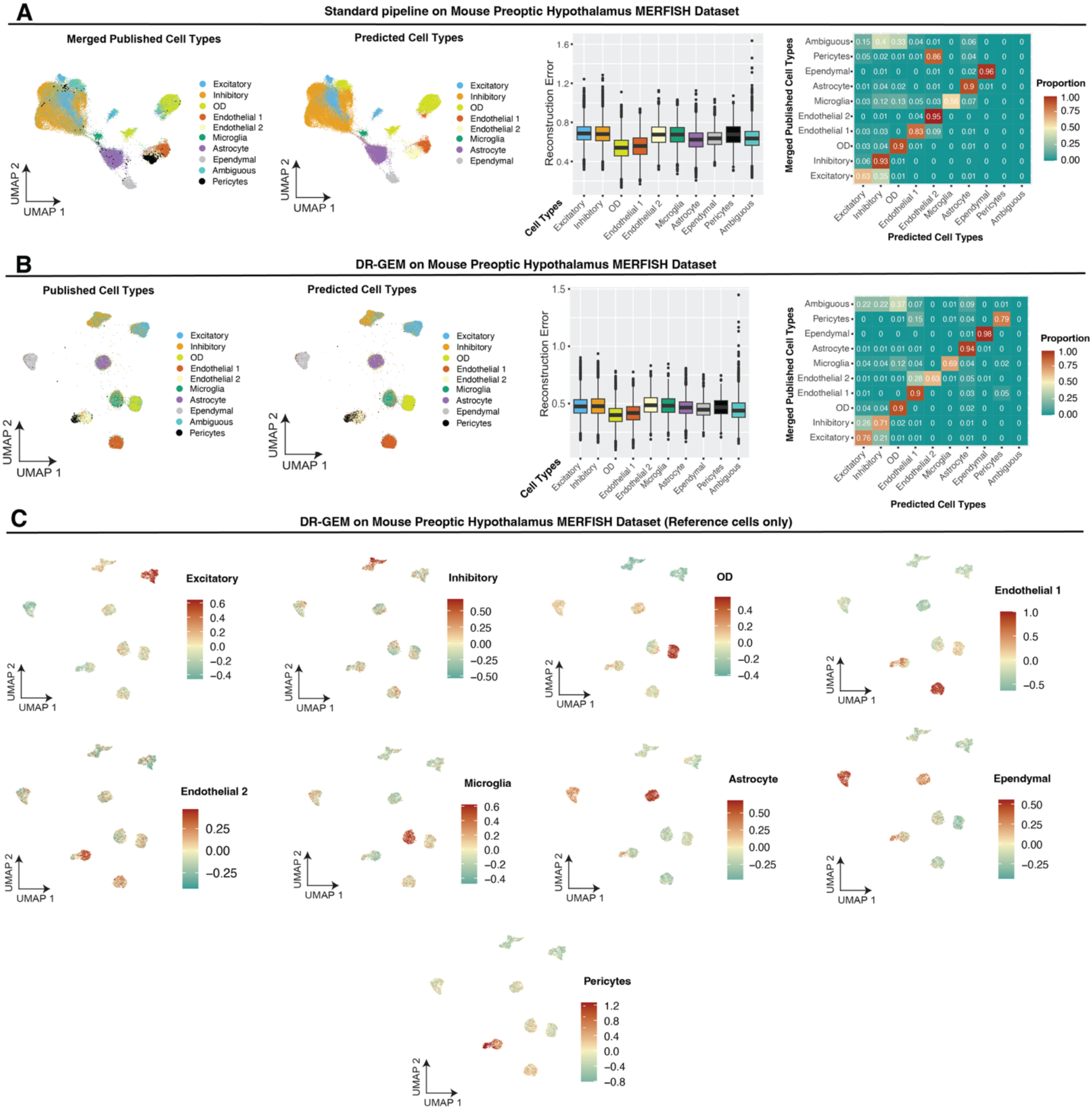
DR-GEM identifies rare cell types and mitigates ambient RNA in a MERFISH dataset profiling the mouse brain. **(A-B)** Performance of standard pipeline **(A)** and DR-GEM **(B)** applied to MERFISH data (Animal 1) of the mouse Preoptic Hypothalamus^14^. UMAP visualization of cells colored by published cell type labels (left) and predicted cell type labels (middle left), boxplots of RE (*y*-axis) in each cell type (*x*-axis; middle right), and proportion of predicted cell type (*x*-axis) in each of the published cell type populations (*y*-axis; right). **(C)** UMAP visualization of high confidence cells in the high confidence reference embedding generated by DR-GEM applied to cells from Animal 1 in a MERFISH dataset of the mouse preoptic hypothalamus. Cells colored by overall expression of cell type signatures derived from a paired scRNA-seq sequencing dataset of the mouse preoptic hypothalamus (**Methods**). **(A-B)** Boxplots middle line: median; box edges: 25^th^ and 75^th^ percentiles; whiskers: most extreme points that do not exceed ± interquartile range (IQR) x 1.5; further outliers are marked individually as black points.

In contrast, with DR-GEM (with “Annotations in the Loop”), Pericytes form a distinct cluster, though they are still close to the Endothelial 2 cells based on Euclidean distance in the PC space and in the UMAP (**Figure 6B-C**). There is no statistically significant difference between RE of Pericytes vs. all other cell types (*p* = 0.45, one-sided Wilcoxon ranksum test) and the range of RE has decreased for the whole dataset. DR-GEM yielded clusters with one-to-one matching to all cell types (**Figure 6B-C**): Excitatory (76% accuracy), Inhibitory (71% accuracy), OD (90%, accuracy), Endothelial 1 (90% accuracy), Endothelial 2 (63% accuracy), Microglia (69% accuracy), Astrocytes (94% accuracy), Ependymal (98% accuracy), and Pericytes (79% accuracy). In addition to the marked improvement in accuracy for Pericytes (from 0% detection under the standard pipeline to 79% accuracy with DR-GEM), DR-GEM outperforms the standard pipeline on all balanced clustering metrics (**Methods, Supplementary Table S1A**). DR-GEM annotations show a more modest alignment with the original annotations for cells originally annotated as Inhibitory neurons and Endothelial 2 cells, potentially indicating that these cells may have been misannotated in the original study. Although cells labeled by the authors as “Ambiguous” were assigned to other cell types, their confidence score is on average lower than scores for all other cell types (*p* = 1.12 × 10^−177^, confidence scores of “Ambiguous” cells vs. all other cell types, one-sided Wilcoxon ranksum test).

### DR-GEM detects and labels transcriptional impact of CRISPR interference in extremely imbalanced Perturb-seq data

Large-scale Perturb-seq datasets, which are starting to become more widely available^47,64,65^, often suffer from extreme class imbalance. In a genome-scale Perturb-seq dataset collected in K562 cultured experimental cells^47^ for example, the number of cells (after filtering for cell quality, **Methods**) harboring each unique genetic perturbation (CRISPR interference) ranges from 2-1,963 whilst the number of high-quality cells with non-targeting controls is 75,305. We applied DR-GEM to this Perturb-seq dataset to test its ability to detect and label the transcriptional impact of a subset of perturbations vs. “core”^47^ non-targeting control (NTC) cells. We selected representative perturbations across a spectrum transcriptional impact magnitude (assessed based on the number of differentially expressed genes (#DEGs), **Methods**), biological functions, and representation in the dataset: *PTPN1* interference (*PTPN1*i, *n* = 371, 0.490% in dataset with NTC, #DEGs = 1,844), MED19i (*n* = 299, 0.395%, #DEGs = 5,528), SRRTi (*n* = 100, 0.133%, #DEGs = 1,799), MRPL424i (*n* = 268, 0.355%, # DEGs =1,245), CASP8AP2i (*n* = 211, 0.279%, #DEGs =1,545), NRDE2i (*n* = 172, 0.228%, #DEGs = 1,091).

In the setting of *PTPN1*i vs. NTC cells in Perturb-seq data, the 2D UMAP visualization of the standard pipeline shows an embedding of cells that form a continuum rather than discrete clusters. Moreover, many of the *PTPN1*i cells are scattered throughout the embedding (**Figure 7A**). In contrast, with DR-GEM, there are two distinct clusters that show a clear 1-to-1 mapping to the genetic perturbation (**Figure 7B-C**), with a tighter range of RE values for both *PTPN1*i and non-targeting control cells (**Figure 7D**). DR-GEM outperforms the standard pipeline across all balanced clustering metrics in both the reference subset (**Supplementary Table S1B**) and across all cells (**Supplementary Table S1A**). Using Phase IV of DR-GEM to assign clusters to perturbation states (**Methods**) shows that DR-GEM (Accuracy = 0.95, Balanced Accuracy = 0.92, Specificity = 0.95, and Sensitivity = 0.89) outperforms the standard pipeline (Accuracy = 0.93, Balanced Accuracy = 0.85, Specificity = 0.93, and Sensitivity = 0.77) in recovering ground truth annotations (**Figure 7E, Extended Figure 2B**).

**Figure 7:**
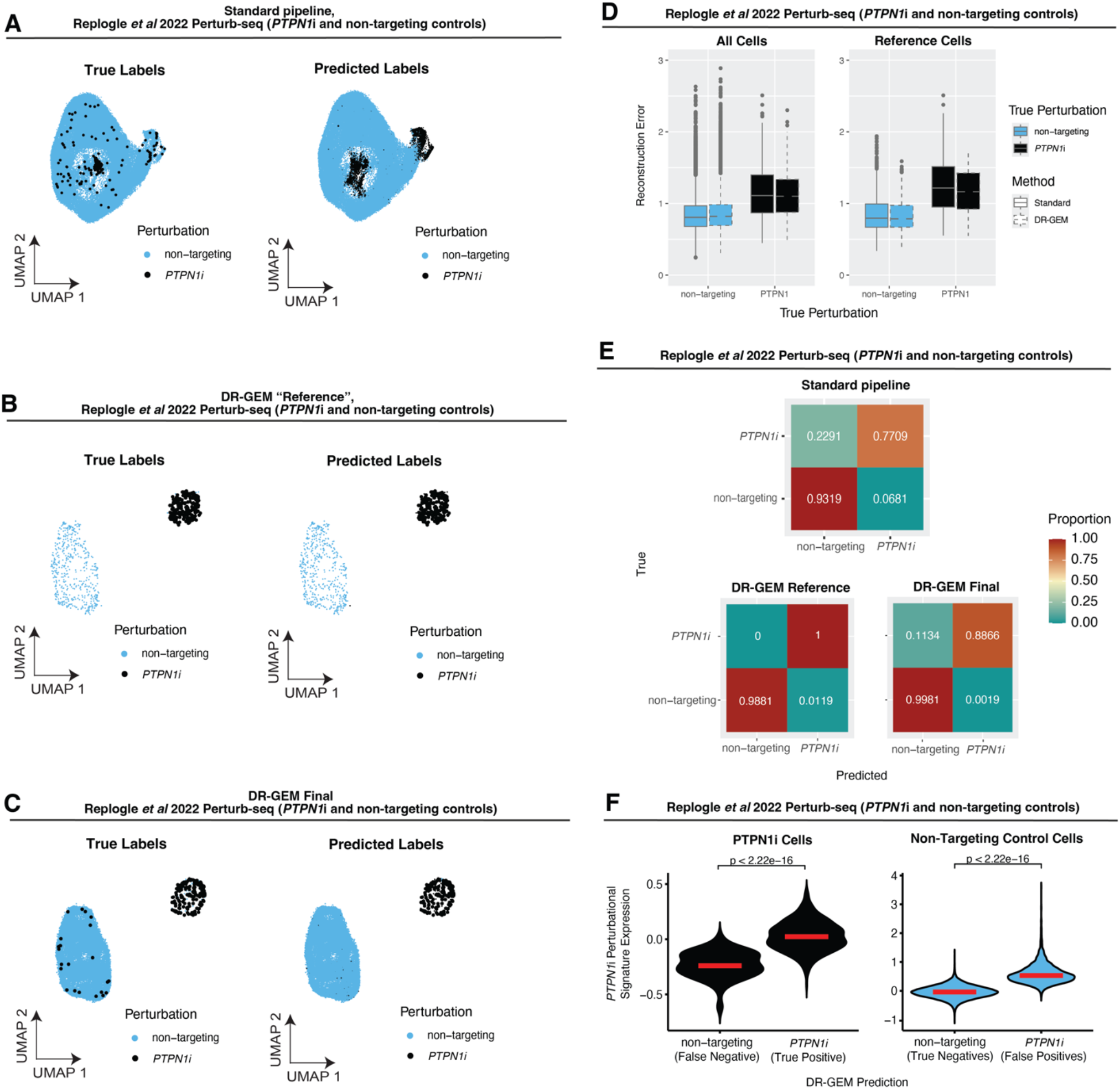
DR-GEM detects and labels transcriptional impact of CRISPR interference in Perturb-seq dataset. **(A-C)** UMAP visualization of cells colored by true perturbation (left) and predicted clusters or perturbations (right) in **(A)** all cells under the standard pipeline, **(B)** high confidence “reference” cells with DR-GEM, and **(C)** all cells with DR-GEM. **(A-C)** Point sizes reflect ground truth annotations based on sgRNA detection (small points represent true non-targeting control cells and large points represent *PTPN1*i cells). **(D)** Boxplots of RE (*y*-axis) in all cells (left) or high-quality reference cells (right), stratified by true perturbation. Boxplots fill color corresponds to true perturbation. Boxplots middle line: median; box edges: 25^th^ and 75^th^ percentiles; whiskers: most extreme points that do not exceed ± interquartile range (IQR) x 1.5; further outliers are marked individually as black points. **(E)** Proportion of predicted perturbations (*x*-axis) in each of the true perturbation groups, as assigned based on sgRNA detection (*y*-axis) when considering (top) all cells under the standard pipeline, (bottom left) high quality “reference” cells with DR-GEM, and (bottom right) all cells with DR-GEM. **(F)** Violin plots of *PTPN1*i perturbational signature expression (*y*-axis) (**Methods**) stratified by DR-GEM prediction (*x*-axis) in *PTPN1*i (left, black) and non-targeting control cells (right, blue). *p-*values derived from one-sided Wilcoxon ranksum test. Red line denotes median of each distribution.

Cells annotated in the Perturb-Seq data as *PTPN1*i are cells in which *PTPN1* single guide RNA (sgRNA) molecules were detected. This annotation may be inaccurate due to ambient sgRNA and because certain cells may express the sgRNA without a functional interference of the target gene expression. In addition, the response of cells to a genetic perturbation may be heterogenous, resulting in visible effects on gene expression only in a subset of the perturbed cells. Using a signature of genes differentially expressed in *PTPN1*i vs. NTC cells shows that cells annotated as *PTPN1*i but predicted by DR-GEM as NTCs also have a lower expression of the *PTPN1*i perturbational signature compared to the other *PTPN1i* cells (p < 2.2 × 10^−16^, one-sided Student’s *t*-test; **Figure 7F**, left). NTC cells predicted as *PTPN1*i cells based on DR-GEM have higher expression of the *PTPN1*i perturbational signature compared to the other NTC cells (*p* < 2.2 × 10^−16^, one-sided Student’s *t*-test; **Figure 7F**, right), suggesting that even in the absence of *PTPN1* genetic interference some cells may exhibit a cell state similar to the *PTPN1*i perturbational state. Importantly, 24% of *PTPN1*i cells have low overall expression (OE < −0.1) of the *PTPN1* perturbational signature, but 59% of these cells are nonetheless predicted by DR-GEM as *PTPN1*i, demonstrating the added value of DR-GEM in detecting perturbation effects beyond standard differential gene expression analyses and signatures.

Similar results were obtained for other genetic perturbations, including *MED19*i (**Figure 8A**), *SRRT*i (**Figure 8B**), *MRPL424*i (**Figure 8C**), *CASP8AP2*i (**Figure 8D**), and *NRDE2*i cells (**Figure 8E**) versus the 75,305 NTC cells from the same genome-scale Perturb-seq dataset^47^. The 2D UMAP visualizations of the standard pipeline shows an embedding of cells that form a continuum rather than discrete clusters. In contrast, the perturbed and NTC cells cluster apart in the DR-GEM reference embedding and in the DR-GEM final embedding. DR-GEM classifications outperform the standard pipeline in the prediction accuracy of the minority/rare perturbed cells (**Figure 8F-G**), Balanced Accuracy and Sensitivity across the whole dataset (**Extended Data Figure 2A**), and in all balanced clustering metrics with both the high confidence reference cells (**Supplementary Table S1B**) and across the whole dataset (**Supplementary Table S1A**). In the cases of *MRPL24*i and *NRDE2*i, DR-GEM detection of the perturbed cells was below 50%, yet false negative cells had a significantly lower expression of the respective perturbation signature compared to the true positive cells (*p* = 9.4 × 10^−6^ for *MRPL24*i cells, *p* < 2.22 × 10^−16^ for *NRDE2*i cells, one-sided Student’s *t*-test; **Extended Data Figure 2B**), indicating that these cells may have not been subject to the perturbation despite the detection of the pertaining sgRNA.

**Figure 8:**
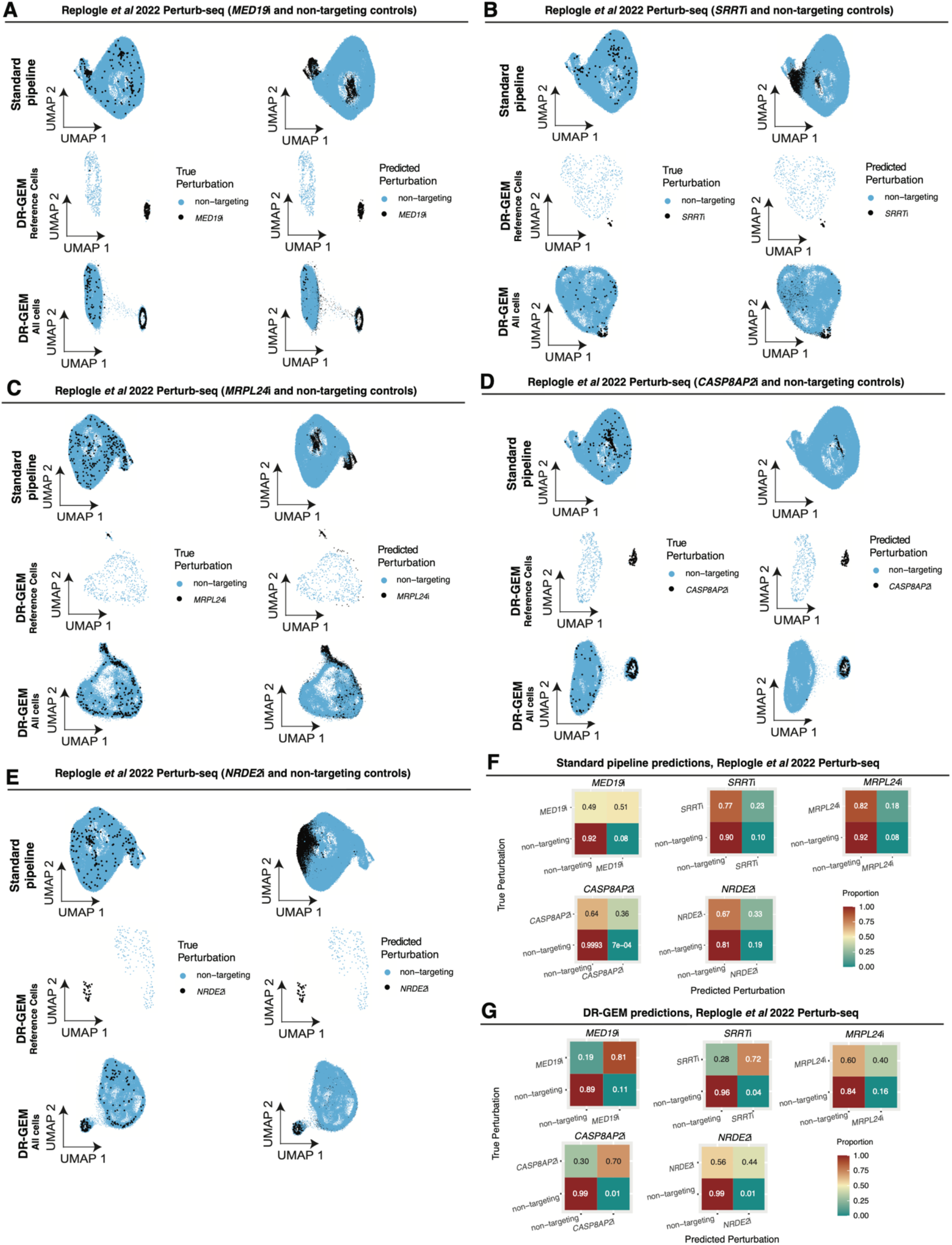
DR-GEM detects and labels genetic perturbational states under extreme class imbalance. **(A-E)** UMAP visualization of cells colored by true perturbation (left) and predicted perturbations (right) in (top) all cells under the standard pipeline, (middle) high quality “reference” cells with DR-GEM, and (bottom) all cells with DR-GEM for non-targeting controls and each of **(A)** *MED19*i, **(B)** *SRRT*i **(C)** *MRPL24*i **(D)** *CASP8AP2*i **(E)** *NRDE2*i. **(F-G)** Proportion of predicted perturbations (*x*-axis) in each of the true perturbation groups, as assigned based on sgRNA detection (*y*-axis), shown for DR-GEM **(F)** and the standard pipeline **(G)**: *MED19*i (top left), *SRRT*i (top middle) *MRPL24*i (top right), *CASP8AP2*i (bottom left), and *NRDE2*i (bottom middle).

These findings demonstrate how DR-GEM can be used to identify cells that were transcriptionally impacted by a genetic perturbation to better annotate Perturb-Seq datasets in an unbiased manner, identify the cell state(s) elicited by genetic perturbations in a manner that is not fully captured by existing supervised (i.e., differential gene expression analyses) and unsupervised analyses (e.g., the standard embeddings and clustering), and determine whether a genetic perturbation is eliciting an already existing or completely novel cell state(s).

## DISCUSSION

Dimensionality reduction and clustering are often first steps in genomics data analysis, providing input to downstream analyses. Here we show that standard pipelines for these data processing tasks are masking unique patterns/entities and do not have mechanisms to reorient the model on high quality data, resulting in suboptimal performances. When applied to single cell data, the most abundant cells and technical noise can dominate the results, skewing the embeddings/clusters, resulting in erroneous annotations, and, at times, leading to total erasure of important biological signals. To address this gap, we developed DR-GEM: a meta-algorithm that incorporates self-supervised data balancing and principles of algorithmic fairness and balanced consensus learning into each component of single-cell annotation pipeline.

DR-GEM consistently outperformed the standard single cell genomics analysis pipeline in faithfully reconstructing latent transcriptional structure, as shown across synthetic benchmarks, spatial transcriptomics and Perturb-seq datasets. The method demonstrated high sensitivity in rare cell recovery (e.g., Mast cells in ovarian cancer, Pericytes in mouse hypothalamus), obtaining reliable embeddings that sharpen biological signal in datasets suffering from ambient RNA, and accurately identified perturbed cell states across extreme class imbalances (e.g., *MED19*i, *SRRT*i, and *PTPN1*i in Perturb-seq data).

DR-GEM achieves this goal by reframing the annotation problem through a self-supervised lens by diagnosing poor embeddings using the RE, upweighting these poorly embedded cells, and leveraging “weak” labels to guide balanced consensus learning that both implicitly upweights samples at risk of underperformance and reorients the model to fit to reproducible signal rather than stochastic noise. It thus takes principles often used in the context of supervised machine learning^68–70^ – where model performances are improved by evaluating model performances and reorienting the optimization function to put more emphasis on samples where the model is still underperforming – and applies these in the context of dimensionality reduction. DR-GEM breaks the feedback loop of bias propagation/amplification while simultaneously generating confidence scores for each annotation with density-normalized subsampling. This confidence-weighted architecture mitigates the impact of artifacts as ambient RNA on the embedding and provides a principled approach to prioritizing ambiguous or low-confidence cells for more extensive consensus validation or exclusion in downstream modeling.

While DR-GEM demonstrates strong performance across diverse datasets and settings, it does have some limitations. The framework adopts a frequentist consensus-based approach, which—while powerful for stabilizing predictions—is more computationally intensive. Future extensions of DR-GEM could benefit from incorporating principles from online learning and adaptive optimization^74–76^, enabling the model to update annotations dynamically as new data become available or as confidence scores start converging. Additionally, while DR-GEM was intentionally designed to be platform-agnostic to maximize generalizability, it currently does not leverage potentially informative platform-specific features such as spatial coordinates in spatial transcriptomics or cross-modality signals in multi-omic datasets. Incorporating these sources of contextual information may further enhance the resolution and specificity of annotations, especially in complex tissue architectures or heterogeneous perturbation screens. We envision that the next generation of distributionally robust self-supervised methods will build upon DR-GEM to incorporate such information in a principled and modular way.

As single-cell datasets continue to grow in size, volume, and complexity, we anticipate that DR-GEM can serve as a foundational approach for single-cell analytics pipelines. DR-GEM is platform-agnostic and compatible with additional domain-specific knowledge via optional gene signature-based labeling, offering flexibility and interpretability in a variety of single-cell settings. Similarly, while here we show DR-GEM performances when applied with PCA it is compatible with other dimensionality reduction algorithms, including non-linear autoencoders and transformers used in single cell foundation models^71–73^ as single cell GPT (scGPT)^72^ and scFoundation^71^, where RE can be used in the same way to reorient the model for more balanced view of biological variation and attention to more diverse biological states.

Beyond single-cell biology, the core methodological principles of DR-GEM are applicable to other settings, and we anticipate further testing and development of DR-GEM for a broader set of applications in the near future. In population-scale datasets like “All of Us”^77^, recent concerns over ancestry visualization using UMAP^78^ underscore how unsupervised embeddings can reinforce biases tied to race and ethnicity, with downstream implications for clinical AI. DR-GEM or its future versions could help mitigate these effects by robustly refocusing learning on minority/rare subgroups. Similar opportunities exist in disease endotyping, such as in asthma^79^ and sepsis^80^, where unsupervised clustering can overlook rare but clinically meaningful subtypes. In psychiatry, multi-omic studies have uncovered biologically distinct subtypes of depression and schizophrenia^81,82^, though such efforts remain vulnerable to class imbalance and noisy ‘omics data. Finally, in digital pathology^77^ and radiomics^,78^, fairness-aware and consensus unsupervised machine learning pipelines may reveal underappreciated morphological patterns in noisy image data. These examples highlight the broader utility of incorporating self-supervision for latent class balancing in machine learning pipelines across biomedical research and biomedicine.

## FIGURES

## TABLES

**Supplementary Table S1:** Summary of (balanced^57^ and standard) clustering performance indices (**Methods**) for **(A)** all cells and **(B)** high confidence “reference” cells, across each dataset tested in this study. Green shading denotes DR-GEM outperforms standard pipeline in given metric for a given dataset; orange denotes vice versa.

**Supplementary Table S2:** Parameter details for ambient RNA simulations in Synthetic Dataset 2. Column A lists all target cell types (cell type A), column B lists all sets of “donor” cell types for each target cell type A: *S*_*Contam,A*_, and column C lists all sets of contamination weights ***w***_*A*_.

## EXTENDED DATA FIGURES

**Extended Data Figure 1:**
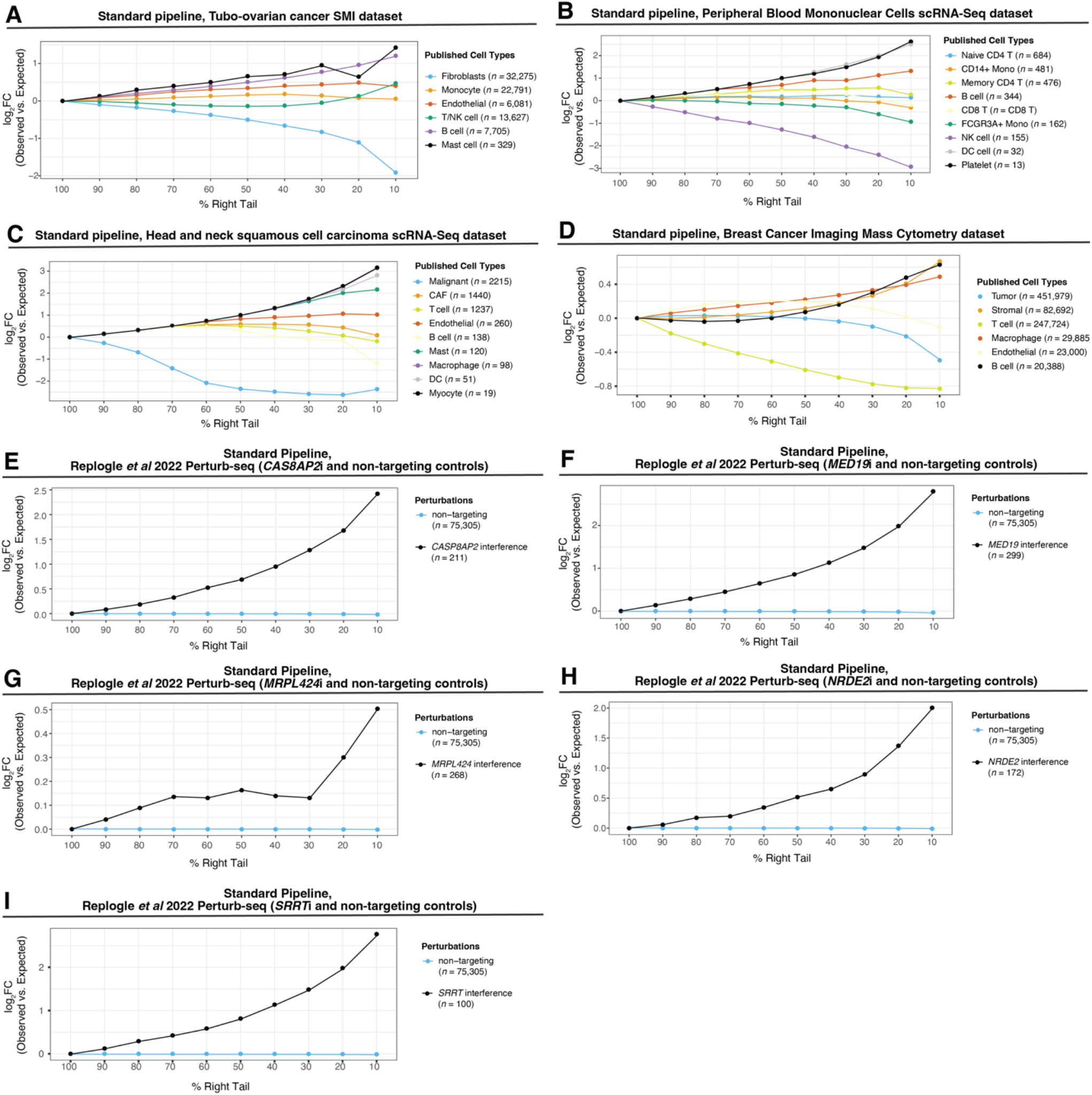
Cells from minority/rare latent classes are more likely to be misrepresented in reduced-dimension embeddings. **(A-I)** Log_2_FC (*y*-axis) of observed percentage of each sample population vs. expected percentage of each sample population (color) in the right-tail (of varying sizes, *x*-axis) of the RE distribution under standard PCA in the following datasets: **(A)** spatial transcriptomics (SMI) profiling of omentum tubo-ovarian tumors**; (B)** scRNA-Seq of peripheral blood mononuclear cells from a healthy donor^50^; **(C)** scRNA-Seq of head and neck squamous cell carcinoma tumors^49^; **(D)** Imaging Mass Cytometry (IMC) profiling of breast cancer tumors^13^; **(E-I)** Perturb-Seq^47^ profiling of non-targeting control cells with **(E)** *CASP8AP2*i cells **(F)** *MED19*i cells **(G)** *MRPL424*i cells **(H)** *NRDE2*i cells **(I)** *SRRT*i cells.

**Extended Data Figure 2:**
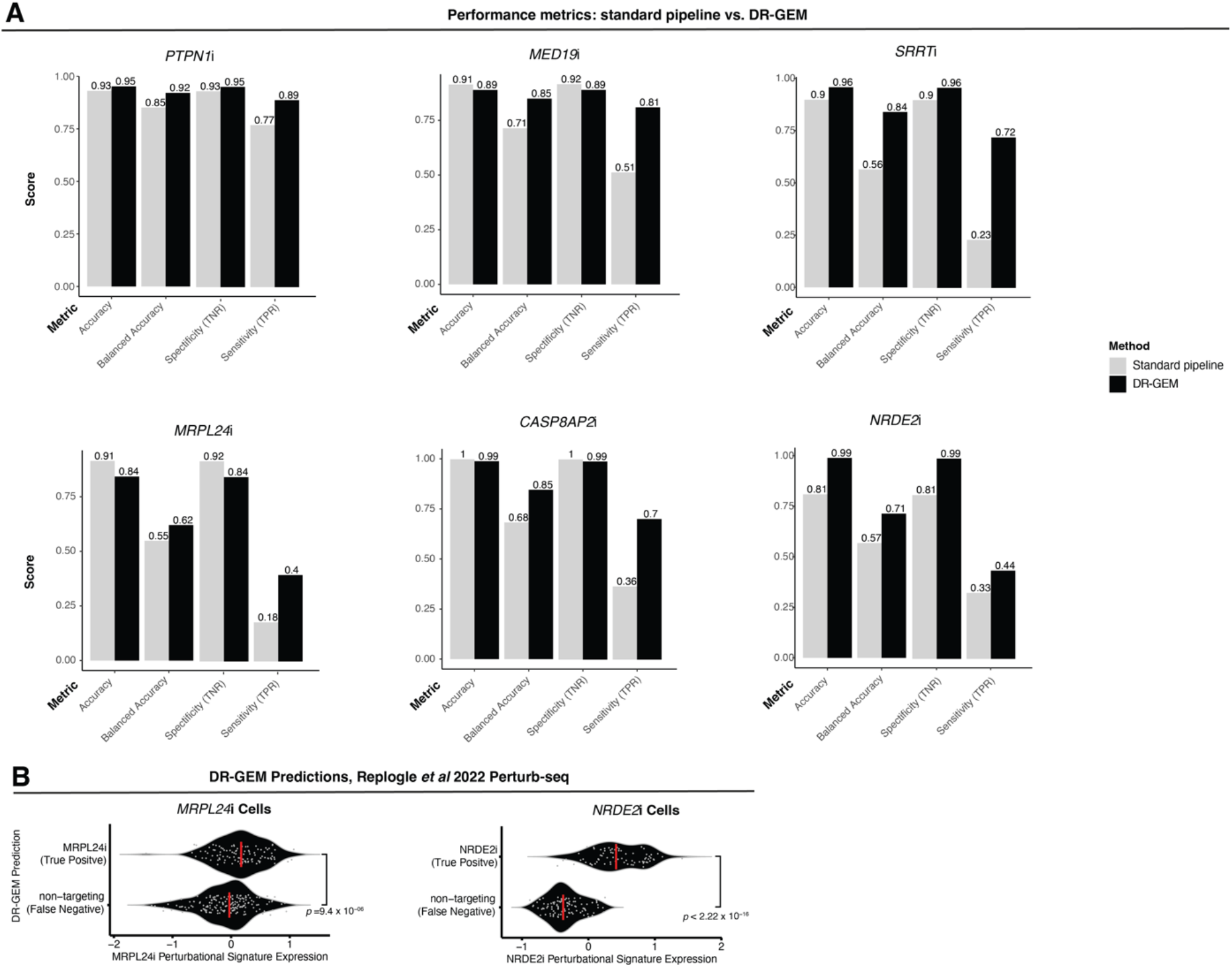
Further performance metrics of Standard pipeline vs. DR-GEM on Perturb-Seq data. **(A)** Barplot of DR-GEM (black) vs. standard pipeline (grey) performance (*y*-axis) across standard classification metrics (*x*-axis) in settings of *PTPN1*i (top left), *MED19*i (top middle), *SRRT*i (top right), *MRPL24*i (bottom left), *CASP8AP2*i (bottom middle), and *NRDE2*i (bottom right) vs. non-targeting control cells. Abbreviations: TNR = true negative rate, TPR = true positive rate. **(B)** Left: Violin plots of *MRPL24*i perturbational signature expression in true *MRPL24*i cells (*x*-axis), stratified by DR-GEM prediction (*y*-axis); Right: Violin plots of *NRDE2*i perturbational signature expression in true MRPL24i cells (*x*-axis), stratified by DR-GEM prediction (*y*-axis); grey points denote each cell, red bar denotes median of each distribution; *p*-values from one-sided Wilcoxon ranksum test.

## DATA AND CODE AVAILABILITY

All synthetic and example datasets in this study are available as tabulated data on Zenodo here: https://zenodo.org/records/15285190. DR-GEM as an end-to-end tool is available as an R package with tutorials to be installed via GitHub (https://github.com/Jerby-Lab/drgem). Please note that all data and code has been prepared for peer-review and may be subject to change. Python-based code for running DRO in PCA as a Linear Autoencoder and Tilted Fair PCA will also available on GitHub (https://github.com/Jerby-Lab/fair-lin-ae) at a later time.

## METHODS

### Synthetic Data Generation

Synthetic single-cell ‘omics datasets were generated using a custom simulation framework. Each dataset consists of count matrices simulating gene expression across *K* discrete “cell types/states”, with parameters selected to reflect common features of real-world data, such as cell type and cell type invariant variation, dropout events, overdispersion, and class imbalance with rare populations.

Gene expression counts were simulated using a two-level negative binomial sampling process. First, for each gene *j* ∈ {1, …, *p*_1_}, and cell type *k* ∈ {1, …, *K*}, a mean expression value was drawn from a negative binomial distribution:

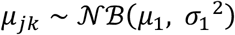

Where *µ*_1_ = 2 and *σ*_1_^2^ = 0.15. In this manner cells of the same cell type overexpress the same marker genes (i.e., genes with a high *µ*_*jk*_). Then, for each cell *i* = 1, …, *N*_*k*_, counts were sampled gene-wise from a negative binomial distribution with cell type specific parameters conditioned on the previously drawn gene-level:

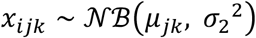

Where *σ*_2_^2^ = 0.5. This process was repeated independently for each cell type *k* ∈ {1, …, *K*}, where *N*_1_, …, *N*_*K*_ cells were sampled from each cell type *k*, respectively (where *N*_1_ + ⋯ + *N*_*K*_ = *N*) yielding a combined count matrix 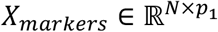 with known cell type labels.

An additional set of *p*_2_ features (where *p*_*2*_ < *p*_*1*,_ and *p* = *p*_*1*_ + *p*_*2*_) of cell-type independent variation was sampled from an analogous two-level negative binomial sampling process. This time having cells randomly assigned to *K*_*2*_ cell states independent of the *K* cell type assignments to generate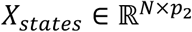.

Concatenating *X*_*markers*_ and *X*_*states*_ yielded counts matrix *X*_*Clean*_ ∈ ℝ^*N* × *p*^.

We simulated negative binomial noise matrix *ε* ∈ ℝ^*N* × *p*^ with *µ*, = 2 and *σ*_3_^2^ = 0.15 such that:

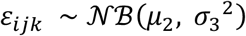

Random dropout was modeled as a binary dropout mask, *D* ∈ {0,1}^*n* × *p*^, generated from a Bernoulli distribution (dropout probability *δ* = 0.05) such that:

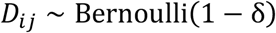

Applying these simulated sources of noise yields a final synthetic counts matrix, *X* ∈ ℝ^*N* × *p*^:

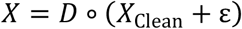

All simulations were performed with a fixed random seed for reproducibility. For subsequent testing and analysis, all simulated datasets were log-normalized and scaled.

### Standard pipeline for single-cell annotations

The standard pipeline for single-cell annotation^50^ comprises three main unsupervised machine learning steps: (1) dimensionality reduction, most commonly via PCA (2) clustering (e.g., SNN clustering) on the first PCs, (3) UMAP visualization, and (4) cluster-based annotation based on marker or signature expression.

Here, we establish some notation for PCA. As is widely known, PCA can be solved analytically via singular value decomposition (SVD) such that for input data *X* ∈ ℝ^*N*×*p*^

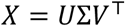

Where *U* ∈ ℝ^*N*×*N*^ contains the left singular vectors, Σ ∈ ℝ^*N*×*p*^is a diagonal matrix of singular values, and V ∈ ℝ^*p*×*p*^ contains the right singular vectors. The lower dimensional embedding *Z* ∈ ℝ^*x*×*m*^ is obtained from the top *m* principal components such that:

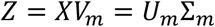

This then yields the rank-*m* approximation (reconstruction) of X:

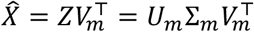

The standard mean squared error (MSE) as RE ***e*** ∈ ℝ^1^ can then be calculated such that for each sample *i*:

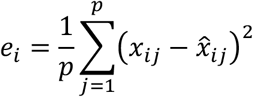

For standard PCA, we used the implementation in the Seurat (v5) R package. The number of PCs retained for each real-world dataset (*m*) was selected using the conventional heuristic of identifying the “elbow point” in the variance-explained plot, representing the inflection at which additional components contribute diminishing returns in explained variance.

### Simulation experiments of synthetic class imbalance

To systematically assess the impact of class imbalance a series of synthetic datasets were simulated with varying levels of class imbalance. Specifically, five discrete cell types (*K* = 5) were simulated with fixed sample sizes for four of the populations (*N*_1_ = 8000, *N*_2_ = 5000, *N*_3_, = 4500, *N*_4_ = 4000) and a variable number of samples for the fifth (rare) cell types:

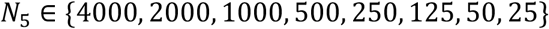

For each of the imbalance settings, 100 independent synthetic datasets were generated with *p*_1_ = 150 and *p*_2_ = 50 and a total of *p* = 200 “genes” using 100 unique random seeds with the two-level generative negative binomial model described in the previous section. Noise (*ε*) and dropouts (*D*) were independently simulated and applied to each dataset.

Each dataset was log-normalized and scaled before application of PCA. RE (***e***) was computed as the mean squared error between the scaled expression data and the reconstruction of the data from the top 10 PCs (*m* = 10). *m* was chosen here based on the “elbow criteria” over 100 datasets where *N*_*5*_ = 4000.

For each simulated dataset and for each cell type, the maximum, mean, and minimum RE was calculated and recorded. Furthermore, the enrichment of each cell type within the top 10% reconstruction error tail (i.e. the top 10% most poorly represented cells) compared to its original baseline proportion was computed with as fold-change:

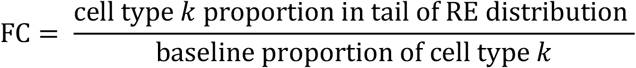

The results are visualized as boxplots stratified by cell type (color) and degree of imbalance (*y*-axis) in **Figure 1B**. Each boxplot summarizes 100 simulations per condition.

### Synthetic Dataset 1

A representative baseline synthetic dataset was utilized as a controlled setting to test different algorithms. This dataset was simulated with the following parameters: *N* = 21,250, *p* = 200, *p*_1_=150, *p*_2_ = 50, *K* = 5, *N*_1_ = 8000, *N*_2_ = 5000, *N*_3_, = 4500, *N*_4_ = 4000, *N*_5_ = 50, *µ*_1_= 2, *σ*_1_^2^ = 0.15, *σ*_2_^2^ = 0.5. A simulated noise matrix (*ε, µ*_2_ = 2, *σ*_3_^2^ = 0.15) and dropout mask (*D*, δ = 0.05) were also applied.

### PCA as Linear Autoencoder

PCA was implemented as a single-layer linear autoencoder (**Figure 2A**) such that it is possible to modify the default loss function and train the model with stochastic gradient descent. It has been previously shown that an autoencoder with a single fully connected hidden layer, a linear activation function and a squared error cost function (RE) trains weights that span the same subspace as the one spanned by the PC loading vector. The loading vectors equivalent to those solved via SVD can be recovered from the autoencoder weights. In other words, for an input data matrix *X* ∈ ℝ^*N*×*p*^, where a *m*-dimensional representation *Z* ∈ ℝ^*N*×*m*^ is obtained (where *m* < *p*) and linear activation weights matrix *W* ∈ ℝ^*p*×*m*^ by minimizing mean-squared error as RE such that:

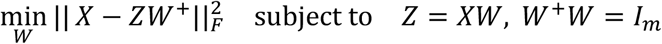

where *W*^+^ is pseudoinverse for *W*. It is equivalent to solving for *V*_*m*_ using the analytical approach for standard PCA.

The implementation of PCA as a linear autoencoder utilized the autoencoder framework in PyTorch^86^ with a single fully connected linear layer of *m*-dimensions. The main training routine receives the model architecture with default initialized weights and a specific loss function (either standard RE or DRO objectives, described next). Models were trained over mini-batches using the Adam optimizer^87,88^ with a fixed weight decay for regularization. Optimization was conducted over user-defined epochs to reach convergence. Model evaluation was conducted from analysis of RE (***e*** ∈ ℝ^1^) as defined above.

### Linear Autoencoder with KL-DRO

In one approach to promote fairness and robustness for PCA, KL-DRO objective was applied for PCA as a Linear Autoencoder.

In this setting the standard RE is defined as:

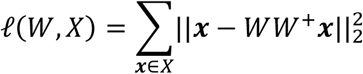

and the KL-DRO objective is defined as:

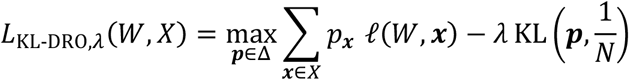

Where KL(·,·) is the Kullback-Leibler divergence between two distributions and *λ* regularization hyperparameter controlling robustness. This objective has been shown to be equivalent to^85^:

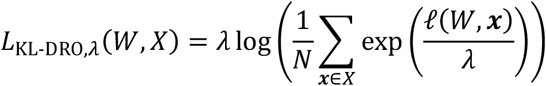

KL-DRO and tilted empirical risk minimization^30^ are mathematically equivalent under a specific transformation of parameters. Both frameworks prioritize worst-case or rare examples by reweighting the empirical loss function in a “tilted” manner (**Figure 2A**).

The hyperparameter *λ* was tuned by testing a set of values in PCA as a Linear Autoencoder:

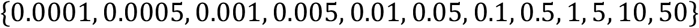

The value *λ* that best minimized maximum RE without a significant cost to overall RE (**Figure 2B**) in the dataset was chosen (*λ* = 0.001).

### Linear Autoencoder with CVaR-DRO

An alternative formulation of DRO is CVaR^51^, a more classic constrained with initial roots in finance and economics. CVaR-DRO objective for PCA as a Linear Autoencoder is as follows:

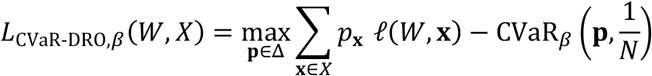

where 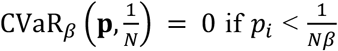, and 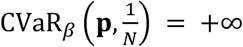 otherwise. The equivalent form here is the optimization of:

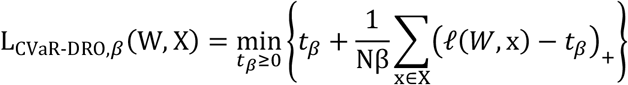

Where *t*_*β*_ is the threshold value that partitions the data to identify the top *β* fraction of reconstruction values. CVaR focuses on minimizing the worst-case (highest) losses — specifically, the average loss in the highest *β*-quantile.

The hyperparameter *β* was tuned by testing a set of values in PCA as a Linear Autoencoder:

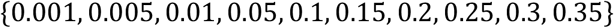

The value *β* that best minimized maximum RE without too much cost to overall RE (**Figure 2B**) in Synthetic Dataset 1 was *β* = 0.01.

### FairPCA

To assess a method where group-wise fairness is enforced in dimensionality reduction, the multiple-objective optimization of FairPCA with tilted reweighting was implemented. This scheme explicitly aims to balances RE across predefined groups. The input dataset *X* ∈ ℝ^*N* × *p*^ is partitioned into its known *K* groups {*X*_1_, *X*_2_, …, *X*_*K*_} where 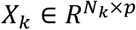 contains all *N*_+_ cells from group *k*. For an orthonormal projection matrix *V*_*m*_ ∈ ℝ^*p*×*m*^ fit from standard PCA, the RE for group *k* is:

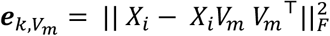

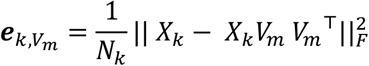

Let 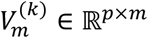 be the projection matrix obtained by performing PCA **only** on group *k*, minimizing RE only within group *k*

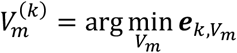

The disparity error for group *k* is then defined as:

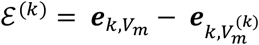

A projection *V*_*m*_ is said to be **fair** if all groups experience the same disparity error:

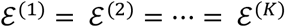

Because exact equality is difficult to achieve in practice, this requirement is relaxed and the strength of Pareto parity^52^ is enforced by the hyper parameter *t* where larger values of *t* force the losses on groups to be as close to identical as possible, whereas lower values of *t* offer the flexibility of reducing the performance gap between groups less aggressively.

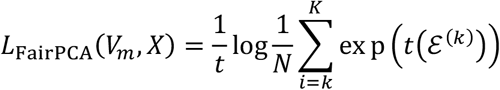

FairPCA models were fit by applying an adapted version of gradient-based multi-objective optimization that has been previously described and implemented^54^. The hyperparameter *t* was tuned by testing the following hyperparameter values:

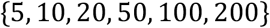

For application on Synthetic Dataset 1, *t =* 10 was sufficient to achieve a level of group-wise parity that successfully minimized the rare cell type RE. The group information was provided to FairPCA, thus providing latent ground truth information to examine performances in favorable conditions that do not recapitulate application in real-world data. However, even in this setting FairPCA failed to identify the rare cells (**Figure 2E**).

### DR-GEM Algorithm

#### Phase I: Distributional Robust Initialization

Let *X* ∈ ℝ^*n*×*p*^ be the normalized and scaled gene expression matrix of *n* cells and *p* genes. DR-GEM begins by fitting a standard unsupervised embedding *E*^(0)^ ∈ ℝ^*n*×*m*^ using a user defined dimensionality reduction method (e.g., PCA), where *m* is user-specified (the “elbow criteria” is recommended). 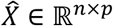 is the reconstructed gene expression matrix from *E*^(0)^. For each cell *i* ∈ 1, …, *n*, a reconstruction error (RE, ***e*** ∈ ℝ^*n*^) is computed as the mean squared error between each row of *X* and 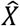. Cells are ranked by RE, and the top *β* percentile (user-defined, default *β* = 0.05) are selected as the “high-risk” subset (*n*^(0)^ = *β* · *n*). A second embedding 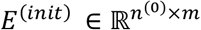 is learned on this subset alone. The remaining cells are projected onto the subspace spanned by *E*^(*init*)^ to yield *E*^(1)^ ∈ ℝ^*n*×*m*^. Clustering (e.g., SNN clustering) is then applied to *E*^(1)^, producing an initial label vector ***c***^(1)^ ∈ {1, …, *K*}^*n*^, referred to as initialized weak cluster labels.

#### Phase II: Balanced Consensus Learning

DR-GEM then fits *r* (user-specified) learners that yield *r* sets of predictions fit on *r* balanced subsamples of the data. The subsample of the dataset is drawn by uniformly sampling from each weak cluster in ***c***^(1)^, thereby balancing the cluster sizes. The uniform sampling procedure is specifically random sampling *min*(*b, N*_*k*_) cells from each cluster *k*, where *b* is user-defined parameter (default *b* = 500) and *N*_+_ is the number of cells in each weak cluster *k*. Let *X*^(2,*r*)^ ⊂ *X* denote a balanced subsample of size *n*^(2,*r*)^, dimensionality reduction projects *X*^(2,*r*)^ to an *m*-dimensional embedding: 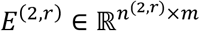, which undergoes subsequent clustering. Because cluster identities can be permuted between runs, labels are aligned to ***c***^(1)^ using the Kuhn–Munkres algorithm (also known as the Hungarian algorithm and implemented as the `lp.assign()` function in the `lpSolve` R package). The Kuhn-Munkres algorithm takes as input the contingency table between two sets of categorical label predictions and determines a one-to-one label mapping. This phase yields an ensemble of cluster predictions for each cell *i*: 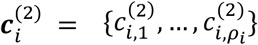 where *ρ*_*i*_ is the number of times a given cell *i* is sampled across the *r* learners.

#### Phase III: Aggregation and Confidence Scoring

The final cluster label ĉ_*i*_ for each cell is assigned via majority vote across the ensemble 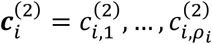 A confidence score *s*_*i*_ ∈ [0,1] is defined as the proportion of iterations in which ĉ_*i*_ was assigned to cell *i* across the *ρ*_*i*_ iterations. A final reference embedding is constructed using a subset of high-confidence cells, defined as the *n*_*t*_ cells with *s*_*i*_ > *t*, where *t* is a user-specified threshold (default: *t* = 0.95). The reference embedding is computed using the same dimensionality reduction method used in previous phases to yield a reference embedding 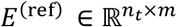. All cells are then projected into this reference subspace to yield embeddings: *E*^(*final*)^ ∈ ℝ^*n*×*m*^ for final visualization and analysis.

#### (Optional) Phase IV: Signature-based Annotation

DR-GEM supports the optional functionality for supervised cluster annotation using curated gene signatures. For any given gene signatures (set of genes) that correspond to markers for a known cell type or state and input gene expression matrix *X* ∈ ℝ^*n* × *p*^ the Overall Expression (OE)^11^ for each cell *i* is computed. Once the OE of all cell types/states of interest are computed across all cells, for each signature OE, one-sided student *t*-tests are performed to test the differential expression of OE values between all possible unique pairs of clusters based on input cluster annotations of the cells. A cluster is assigned to a specific cell type if the cells assigned to it have a significantly higher OE (defined by *z*-score from one-sided *t*-tests, where *z*-score > 10) of the cell type signature compared to all other clusters. Clusters with overexpression of more than one cell type signatures by this definition undergo a tie-breaking scheme utilizing the differential OE effect sizes: the cluster is assigned to the cell type with the highest OE *z*-score compared to other clusters. The final output is a set of cell type or state labels mapped from input cluster annotations.

#### Parameter Selection for DR-GEM

Throughout the study clustering was performed via the shared nearest neighbor (SNN) modularity optimization-based clustering algorithm implemented in the <monospace>Seurat </monospace>R package (v5). SNN first calculates

*k*-nearest neighbors to construct a SNN graph and applies a modularity function to determine clusters. All parameters were set to the default, except for the clustering resolution. For each dataset, a set of resolution values were tested under the standard pipeline; the resolution value that best optimized Adjusted Mutual Information (AMI) between ground truth labels and clusters was selected.

Visualization on a 2-dimensional UMAP was performed with all default parameters within <monospace>Seurat </monospace>R package (v5) for each dataset. In instances that called for the projection of cells to a reference embedding, the reciprocal PCA (RPCA) is used to derive reciprocal PCs that span the target subspace. The implementation within the <monospace>Seurat </monospace>R package was used. The number of PCs used is equal to the value determined at the beginning of each pipeline for each dataset tested using the “elbow criteria”.

#### Clustering performance metrics

Clustering performances were quantitatively evaluated using both standard metrics and analogous metrics that account for class imbalance^57^. The standard metrics used are the Adjusted Rand Index (ARI), Adjusted Mutual Information (AMI), Homogeneity Score, Completeness Score, and V-measure as implemented in the <monospace>aricode </monospace>(v1.0.3) R package. Since these metrics do not account for unequal class prevalence, a balanced version of each of these metrics (bARI, bAMI, bHomogeneity, completeness, and bV-measure, where “b” stands for balanced) recently published^57^ and available at https://github.com/hsmaan/balanced-clustering was used. These balance metrics reweight contributions from each ground truth class equally – mitigating the dominance of majority classes in the evaluation metrics. These metrics were computed for each test dataset to benchmark DR-GEM compared to the standard pipeline (**Supplementary Tables S1A-B**).

#### Synthetic Dataset 2: ambient RNA simulation

To simulate contamination in synthetic single-cell data, a controlled signal mixing across specific cell types was added. First, gene expression matrix *X*_clean_ is simulated. For each of cell type A, a set *S*_*Contam,A*_ of contaminating cell types is defined as well as a set of contamination weights (***w***_***A***_, between 0 and 0.6). Then, for each cell *i* of cell type A, a contamination vector is defined as:

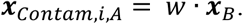

Where *w* is a contamination weight randomly sampled from ***w*** (i.e. *w* ∈ ***w***_***A***_) and ***x***_***B***_ is a single cell gene expression vector of cell type B (where *B* ∈ *S*_*Contam,A*_) sampled from *X*_clea*n*_. Repeating this procedure for all cells *i* ∈ {1, … *n*} of all cell types *k* ∈ {1, …, *K*} yields a contamination matrix (*X*_*Contam*_ ∈ ℝ^*n* × *p*^) that is added to target cells: *X* _*Contam*_ + *X*_clea*n*_ to yield a counts matrix. The specific parameters used to generate contamination in Synthetic Dataset 2 are provided in **Supplementary Table S2**. After introducing contamination, noise and technical dropouts were simulated as described above. After applying noise (*ε*) and drop out (*D*) as in Synthetic Dataset 1, this gene count matrix was log-normalized and scaled for subsequent testing and analysis.

#### Data processing for tubo-ovarian cancer SMI dataset

The SMI dataset of the Discovery cohort from a previous publication^11^ was downloaded as the file: “ST_Discovery.rds” from Zenodo at (https://zenodo.org/records/12613839). For testing the default DR-GEM algorithm, the data was filtered to correspond to only the non-malignant cells from omentum tumors (**Figure 5A-C**); the data subset consisted of 82,808 cells profiled from 47 omentum tumors that were sampled from 41 patients. The raw counts matrix was log-normalized and scaled for subsequent testing.

For testing the DR-GEM with “Annotations in the Loop”, the dataset was used with all cells retained (**Figure 5D, F**), corresponding to a total of 491,792 cells from 100 tumors (53 adnexal, 47 Omentum) sampled from 58 patients. The raw counts matrix was log-normalized and scaled for subsequent testing.

The final “broad” cell type labels previously defined in the original paper were used as ground truth for both settings.

#### Data processing for tubo-ovarian cancer MERFISH dataset

The MERFISH dataset of the Validation 2 cohort from a previous publication^11^ was downloaded as the file “ST_Validation2.rds” from Zenodo at (https://zenodo.org/records/12613839). For testing the DR-GEM with “Annotations in the Loop”, the dataset was used with all cells retained (**Figure 5E, G**), corresponding to a total of 425, 578 cells from 4 adnexal tumors each sampled from 4 patients. The raw counts matrix was log-normalized and scaled for subsequent testing. The final cell type labels previously defined in the original paper were used as ground truth.

#### Data processing for the mouse preoptic hypothalamus MERFISH dataset

The MERFISH dataset was downloaded from Dryad (https://datadryad.org/dataset/doi:10.5061/dryad.8t8s248). The data from Animal 1 was used, consisting of 491,792 cells and 155 genes. From the raw counts matrix, “transcripts per million” (TPM) values were derived as previously described^14^. The TPM matrix was then scaled before subsequent analysis and testing. Pairwise one-sided *t*-tests were applied to determine the number of DEGs between all unique pairs of the most granular cell subtype labels provided in the original paper. The criteria for two cell subtypes to be kept separate is if the two subtypes have more than 30 DEGs (defined as Benjamini Hochberg (BH) corrected *p*-value < 0.01 and |log_2_FC| > 2). The only cell subtypes that were kept separate were the endothelial subtypes. Endothelial 2 and Endothelial 3 cells were merged into “Endothelial 2”, whilst “Endothelial 1” was kept separated.

#### Perturb-seq data processing

A Perturb-seq dataset of a CRISPR interference screen in K562 leukemia cells was downloaded as raw counts data made available upon request from the original authors. With consideration for data quality, only control samples that corresponded to the “core” control cells (*n* = 75,305) as defined in the original paper^47^ were used. For cells that were perturbed, 334 perturbations with strong transcriptional phenotype (# DEGs > 1000, as defined by the mean number of genes determined as significantly differentially expressed between perturbed and control cells via the Anderson-Darling test and Wilcoxon ranksum test), with at least 25 cells, and more than 30% knockdown were considered. From this set of filtered perturbations, 6 perturbations (*PTPN1*i, *MED19*i, *SRRT*i, *MRPL424*i, *CASP8AP2*i and *NRDE2*i) were selected to represent a range of transcriptional impact, biological functions, and frequency in the dataset.

#### DR-GEM with “Annotations-in-the-loop”

DR-GEM with “Annotations-in-the-loop” is Phase I-III of the DR-GEM algorithm, but with Phase IV utilized in Phase I and II. In this variant of DR-GEM, the weak cluster labels at the end of Phase I: ***c***^(1)^ ∈ {1, …, *K*}^*n*^, are mapped to cell type or state labels using the optional Phase IV of DR-GEM. In Phase II, of DR-GEM with “Annotations-in-the-loop”, clusters derived in each *r* iteration are mapped to cell type/state labels, again using the optional Phase IV of DR-GEM. In this version of DR-GEM, the scores and final predictions are determined from the ensembles of cell type/state labels in Phase II.

## CONTRIBUTIONS

C.Y.Y. and L.J., designed the study. C.Y.Y. conducted the research and developed the main software.

C.Y.Y and L.J interpreted the results. M.W.S. supported development of simulation framework and software. D.Z. supported software development for implementing DRO. C.Y.Y. and L.J. wrote the manuscript. L.J. obtained the funding and supervised the study. All authors reviewed and approved the manuscript.

## ACKNOWLEDGEMENTS

We thank Subin Kim for support on preliminary data processing used in earlier versions of this work; Shiori Sagawa and John J. Cherian for insightful conversations on distributional robust optimization. C.Y.Y. is supported by the Stanford Graduate Fellowship in Science & Engineering and the Stanford Medical Scientist Training Program. M.W.S. is supported by the National Institutes of Health (NIH) T15 LM007033 and Stanford Data Science. L.J. is a Chan Zuckerberg Biohub Investigator and an Allen Distinguished Investigator. L.J. holds a Career Award at the Scientific Interface from the Burroughs Wellcome Fund and a Liz Tilberis Early Career Award from the Ovarian Cancer Research Alliance (OCRA). This study was supported in part by the BWF (1019508.01; L.J.), OCRA (889076; L.J.), National Institutes of Health (NIH; U01HG012069; L.J.), and funds from Chan Zuckerberg Biohub (L.J.).

## Notes

https://github.com/Jerby-Lab/drgem

https://zenodo.org/records/15285190

### Competing Interest Statement

The authors have declared no competing interest.

